# A dual sensor regulates P-glycoprotein’s structural plasticity

**DOI:** 10.64898/2026.03.13.711505

**Authors:** Michael Kamel, Jan-Hannes Schäfer, Valeria Jaramillo-Martinez, Nghi N. B. Tran, Devin L. Mangold, Dmitry Shvarev, Kilian Schnelle, Kristian Parey, Dovile Januliene, Ina L. Urbatsch, Arne Moeller

## Abstract

P-glycoprotein is an efflux pump with an exceptionally broad substrate profile which drives its profound clinical impact. Despite its biological importance, it remains obscure how P-glycoprotein achieves its polyspecificity and how substrate binding and the lipid environment stimulate its activity. Structural data highlight the importance of transmembrane helices 4 and 10, which surround the binding pocket, and identify them as key players in substrate recognition.

Here, we used cryo-EM to study P-glycoprotein in detergent and nanodiscs to strategically leverage environment- and substrate-dependent phenotypes. This approach allowed us to decipher unexpected and distinct roles of transmembrane helices 4 and 10, which structurally explain differences in ATPase activity. Our data highlights helix 4 as an environment sensor and helix 10 as the key player in substrate recognition constituting a dual regulation mechanism for the functional plasticity of P-glycoprotein, and visualizes the intricate interplay between a membrane protein and its environment.

## Introduction

P-glycoprotein (P-gp or ABCB1, MDR1) is a well-investigated ATP-binding cassette (ABC) transporter and a polyspecific efflux pump. High expression levels of P-gp have been identified in the small intestine, liver, kidneys, and blood-brain barrier, promoting rapid detoxification of absorption-sensitive or absorption-restricted tissues. P-gp exports many structurally unrelated amphiphilic and either neutral or cationic compounds ^1^, often used in clinics as drugs, thereby reducing drug absorption and limiting their effective clinical use. Accordingly, P-gp overexpression in cancer cells is linked to the onset of multidrug resistance (MDR), allowing tumor cells to export and thus evade many established chemotherapeutics ^2^.

P-gp is expressed as a single polypeptide chain, folded into two transmembrane domains (TMDs) and two highly conserved nucleotide-binding domains (NBDs), with the two pseudosymmetric halves connected by a flexible linker. Based on the TMD fold, it is classified as a type IV ABC transporter and belongs to the B-family of mammalian ABC transporters ^3,4^. For the ABC transporter superfamily, substrate translocation is generally understood by the alternating-access model ^5^, in which ATP binding, hydrolysis, and the release of ADP and inorganic phosphate fuel transitions between the inward-facing (IF) and outward-facing (OF) conformations ^6^ to facilitate transport. In the IF conformations, the NBDs are separated, and the intracellular gate of the transporter is open, allowing access of substrates to a large hydrophobic binding pocket located between the two TMDs. ATP binding induces dimerization of the NBDs, leading to a significant reorganization of the TMDs into OF conformations and substrate release. In general, transport substrates stimulate the ATPase activity, indicating cross-communication between the catalytic and transport domains ^7^.

Commonly, the two central transmembrane helices 4 and 10 (TM4 and TM10), lining the drug-binding pocket, exhibit pronounced flexibility. They can bend in response to substrates and inhibitors, highlighting their importance as key players in substrate recognition ^8–10^. Besides the central drug-binding pocket, an additional inhibitor-binding site (vestibule) was identified between the proposed access tunnel and the central-binding pocket, underscoring potential differences between substrates and inhibitors ^10^. Despite multiple P-gp structures in the presence of various compounds, the conformation of the drug-free state remains controversial, likely due to the transporter’s high flexibility in its apo state ^11,12^. Interestingly, a recent study resolved an apo structure of human P-gp in saposin nanoparticles in a new IF asymmetric conformation ^8^. However, it is not yet clear how and at which point this conformation can be incorporated into P-gp’s transport cycle.

P-gp exhibits elevated basal and drug-stimulated ATPase activity in nanodiscs compared to detergent micelles ^13,14^, which is also the case for many other ABC transporters ^15,16^. Moreover, not only activity, but also conformation can be environment-dependent, as demonstrated recently for the bacterial ABC transporter MsbA ^17,18^. We strategically employed different hydrophobic environments to capture and characterize multiple apo and verapamil-bound conformations of human P-gp. While human and mouse P-gp display high sequence identity (87%), they exhibit different substrate specificities ^19–21^, therefore, we additionally explored interspecies divergence by complementing our data on human P-gp with cryo-EM structures of detergent-stabilized mouse P-gp in apo and verapamil-bound states. Altogether, in this study, we probed effects of the membrane environment, substrate binding, and species-specific variation on the transporter structure and dynamics. By uncovering intermediate states, our results highlight distinct roles of TM4 and TM10 in substrate recognition and transport. TM4 acts as environment sensor, while TM10 responds to verapamil. Overall, a model emerges that explains how P-gp senses changes in the membrane and the presence of substrates, which serve to explain variations in activity.

## Results

### Cryo-EM of P-gp in the apo state

First, we determined cryo-EM structures of detergent-solubilized (LMNG/CHS) human and mouse P-gp in the apo state. Cholesterol hemisuccinate (CHS) was added during purification, as it has previously been shown to stimulate P-gp ATPase activity (Figure S1A and C) ^14,22^. Under these conditions, for human P-gp, we resolved three different IF conformations, termed IF-wide^+^, IF-wide, and IF-narrow (Figure 1A, B, and C), based on the degree of NBD separation, reflected by the distance between the C*α* atoms of the catalytic glutamates (E556 – E1201, Figure S2). 48% of the particles occupied the IF-wide^+^ conformation, displaying the widest NBD separation (E556 – E1201 distance of 44 Å), 30% populated the IF-wide (36 Å), and 22% the IF-narrow (31 Å) (Figure 1 and S2). Interestingly, for mouse P-gp, only one major IF-wide^+^ conformation (Figure 1D) was resolved with an inter-catalytic glutamate distance (E552-E1197) of approximately 41 Å. Thus, in the absence of substrate, both transporters favor wider conformations in LMNG/CHS, while human P-gp appears to additionally exert greater flexibility than its mouse counterpart, as further supported by the 3D variability analysis (Figure S3).

**Figure 1.**
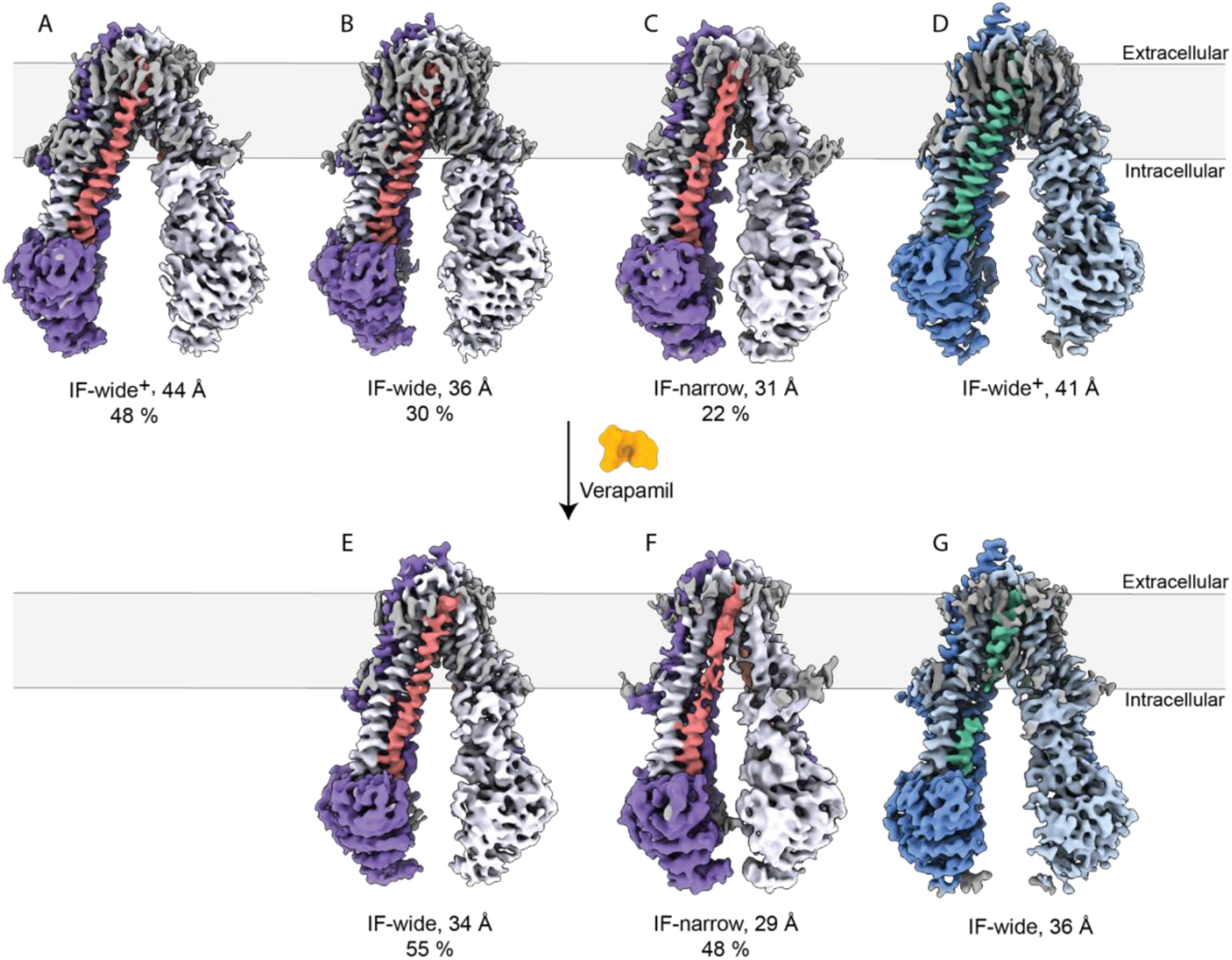
Cryo-EM analysis of P-gp in detergent micelles. Cryo–EM structures of human (white and purple) and mouse P-gp (blue and light blue) in the drug-free and verapamil-bound states in detergent micelles (LMNG/CHS). The measured distance between the catalytic glutamates (E556 and E1201 for human P-gp, E522-E1197 for mouse P-gp) is indicated for each conformation. The TM10 helices are colored in coral and green, respectively.

In all densities, we detect multiple CHS molecules that wrap around the protein, forming a belt around the upper half of the transmembrane domain (TMD), thus indicating how CHS increases the thermal stability of the protein and thereby possibly stimulates the ATPase activity of P-gp (Figure S4)^14,23–25^. Additionally, an LMNG molecule resides in the substrate-binding pocket in the IF-wide conformation of human P-gp (Figure S5A). This is in line with previous observations describing detergents as transport substrates and modulators for P-gp, which can inhibit its ATPase activity at higher concentrations ^26–28^.

For cryo-EM analyses in nanodiscs, detergent-purified human P-gp was reconstituted into the respective belt protein, carrying brain polar lipids and cholesterol (80:20 w/w), following established protocols with minor adjustments (Figure S1D and E)^10,29^. We first employed the largest membrane scaffold protein MSP2N2, which allowed sampling of the natural conformations of the bacterial ABC transporter MsbA ^17^. For human P-gp, in the apo-state, a single major IF-narrow conformation was resolved with an inter-catalytic glutamate distance of 30 Å (Figure 2A). Remarkably, in the smaller MSP1D1 nanodiscs, only 14% of the particles exhibited such an IF-narrow conformation (Figure 2B), which could only be poorly resolved due to the low abundance. The remaining particles populated an asymmetric conformation (IF-asym), which cannot be categorized as a typical IF conformation (Figure 2C) and resembles the recently resolved structure of P-gp in saposin nanoparticles ^8^. Curiously, this IF-asym was neither detected in LMNG/CHS micelles nor in MSP2N2 nanodiscs.

**Figure 2.**
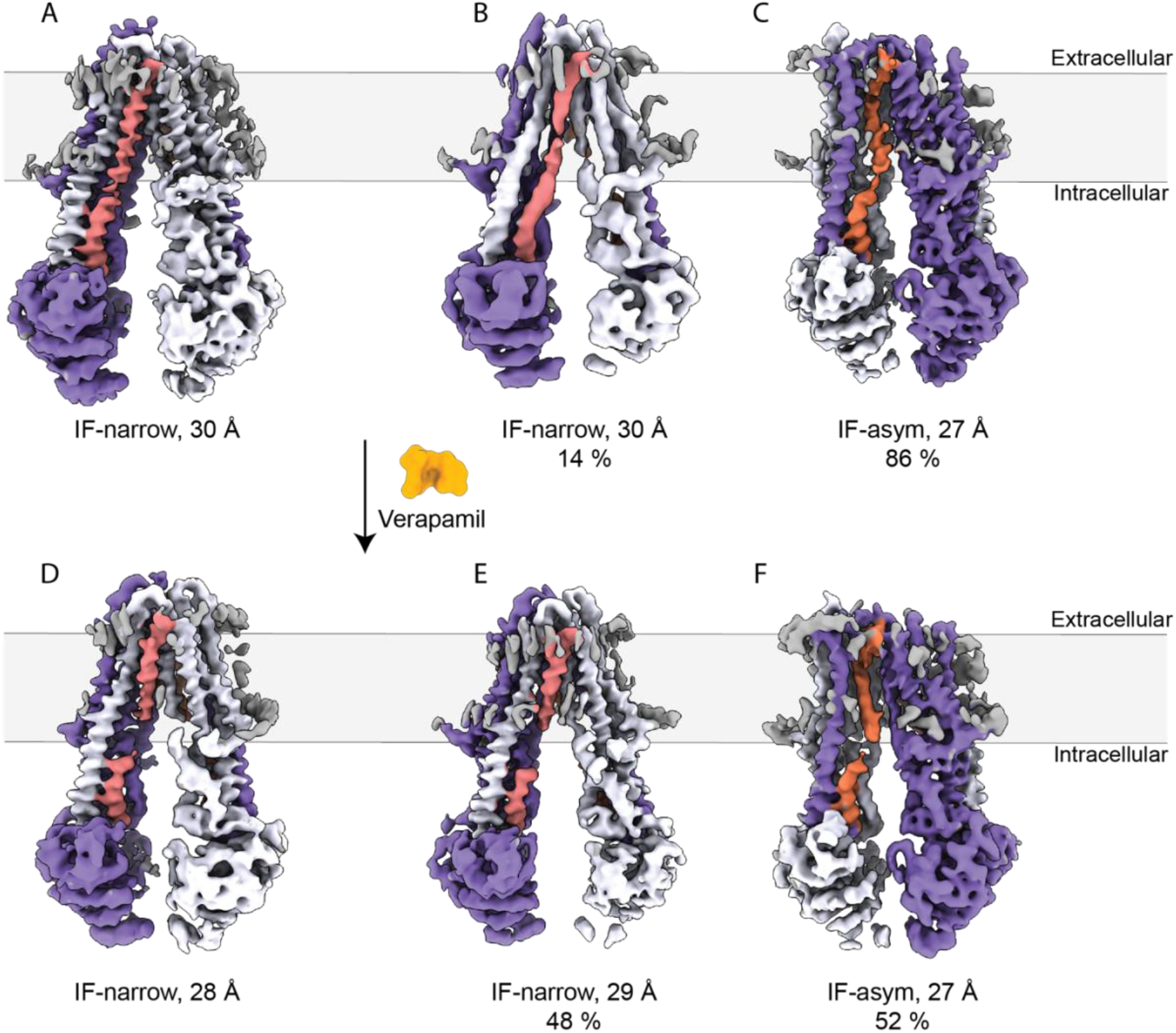
Cryo-EM analysis of P-gp in nanodiscs. Cryo–EM structures of human (white and purple) P-gp in MSP2N2 nanodiscs in apo (A) and verapamil-bound states (D). Cryo-EM structures of human P-gp in MSP1D1 in apo (B and C) and verapamil-bound states (E and F). The TM4 helices are orange and TM10 are in coral.

### Cryo-EM of verapamil-bound P-gp

Verapamil is a calcium channel blocker of broad clinical significance for the treatment of hypertension, angina, and arrhythmias ^30^. It is a well-studied substrate for P-gp and has been described as an efflux modulator ^31,32^, which is known to stimulate the ATPase activity of P-gp. Therefore, we initially measured and compared the activity of human and mouse P-gp using a colorimetric ATPase assay over a range of verapamil concentrations from 0 to 1 mM in detergent micelles and nanodiscs. Both human and mouse counterparts showed comparably low basal ATPase activity (7 and 25 nmol/min/mg, respectively) in detergent micelles, with verapamil stimulating mouse P-gp approximately two-fold more potently than its human counterpart (Vmax of 627 and 290 nmol/min/mg, respectively) (Figure 3A). When reconstituted in nanodiscs, the human transporter exhibited an elevated basal ATPase activity (135 and 277 nmol/mg/min in MSP1D1 and MSP2N2, respectively), which was further stimulated by verapamil (Vmax of 270 and 604 nmol/min/mg), consistent with previous reports (Figure 3B)^10^. These observed differences may reflect intrinsic, species-specific properties of P-gp and arise from distinct ligand-binding modes or conformational states.

**Figure 3.**
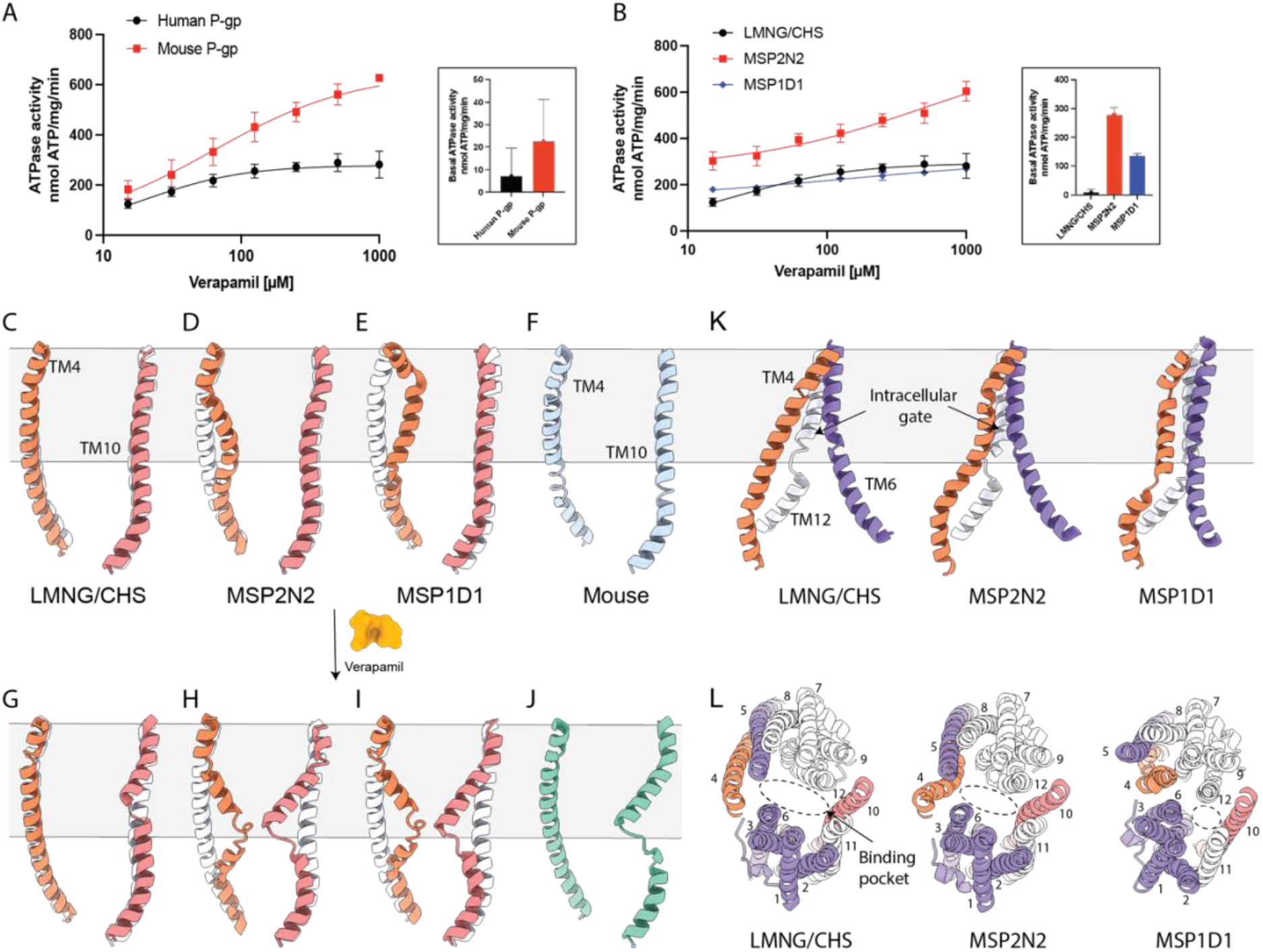
Plasticity of TM4 and TM10. A) ATPase activity of human and mouse P-gp in LMNG/CHS micelles. B) ATPase activity of human P-gp in the different hydrophobic environments used in this study. C) In LMNG/CHS, both TM4 (orange) and TM10 (coral) helices of human Pgp remain straight in IF-narrow and IF-wide conformations, the latter is shown as a guide (white helices). D) The IF-narrow conformation in MSP2N2 nanodiscs shows pronounced bending in TM4 along with modest bending in TM10. E) The IF-asym conformation in MSP1D1 shows strong kinking in TM4 and a straight TM10. F) Similar to human P-gp, the mouse transporter shows straight helices in the apo state in LMNG/CHS. G) In human Pgp, verapamil induces modest bending of TM10 in the IF-narrow conformation in LMNG/CHS, whereas H) in MSP2N2, both TM4 and TM10 are significantly bent. I) A double bent conformation is also observed in the IF-narrow conformation in the small nanodiscs MSP1D1 in the presence of verapamil. J) Verapamil induces significant bending of TM10 in mouse P-gp in LMNG/CHS. K) In contrast to the open intracellular gate of human Pgp in LMNG/CHS, the access of the intracellular gate is dramatically reduced in MSP2N2 and in MSP1D1. L) The bending of TM4 and TM10 reshapes the substrate-binding pocket in the different hydrophobic environments: the cavity (dotted oval) is the widest in detergent micelles, narrows in MSP2N2, and collapses in the IF-asym conformation in MSP1D1

Next, we determined the cryo-EM structures of the verapamil-bound states of human and mouse P-gp in detergent micelles. Notably, in LMNG/CHS, upon verapamil binding to the human transporter, the fraction of the IF-narrow conformation increased from 22% to 48% (Figure 1F). Moreover, the IF-wide^+^ conformation was not observed in the verapamil-bound state. For mouse P-gp, an IF-wide conformation (Figure 1G) was resolved with a 36 Å separation between the NBDs, thus also narrower than the apo state (∼41 Å).

In analogy to the apo state, we resolved a single IF-narrow conformation in MSP2N2 nanodiscs (∼28 Å) (Figure 2D). Intriguingly, in MSP1D1 nanodisc, IF-narrow and IF-asym conformations, like in the apo state, are still present. Nevertheless, their distributions changed, as the fraction of IF-narrow conformations (∼29 Å) increased from 14% to 48% (Figure 2E and F). Altogether, our data show that the conformational spectrum remains similar in the apo and substrate-bound states. However, substrate binding shifts the particle distribution toward narrower conformations across all tested hydrophobic environments.

### Conformational changes in TM4 and TM10

Plasticity of the transmembrane helices TM4 and TM10 is important for substrate binding and transport in P-gp ^10,29,33–35^. Our comparisons between detergent-and nanodisc-stabilized P-gp show that not only substrate binding but also the environment itself has a substantial impact on these helices.

In detergent, all IF conformations (IF-wide^+^, IF-wide, or IF-narrow) of human P-gp display straight TM4 and TM10 helices in the apo state (Figure 3C). However, in MSP2N2 nanodiscs, TM4 bends inwards toward the central cavity between residues P223 and E243, shifting the helix by ∼10 Å, as measured at residue W232, representing a pronounced displacement of TM4 (Figure 3D, orange helix). In contrast, TM10 of the same structure is only slightly displaced between residues P866 and D886 (Figure 3D, coral).

In the smaller MSP1D1 nanodiscs, the sparsely populated IF-narrow conformation resembles the one resolved in MSP2N2 nanodiscs. In contrast, the dominating IF-asym conformation exhibits a different and unusual conformation. Here, TM4 exposes an alternative configuration, the helix breaks at two main points, S222 and E243, which divide it into three segments (Figure 3E, orange helix). This brings T238 in proximity to TM6 and residue S222 to cross over to TM12 on the pseudo-symmetric half of the transporter. This leads to sealing of the intracellular gate (Figure 3K) and causes the binding pocket to collapse (Figure 3L), likely hindering substrate binding, consistent with previous observations ^8^. Such a configuration promotes numerous rearrangements in other TMDs, which are most pronounced in the outward movement of TM5 to accommodate TM4 kinking. Similarly, an outward movement of TM1/TM2 and TM7/TM8 is also observed at the level of the outer leaflet of the membrane.

Generally, movements in TM4 and TM10 reshape the substrate-binding pocket, altering the accessible volume for substrate binding. The volume is the largest in detergent micelles, where TM4 and TM10 are straight, decreases in the MSP2N2 nanodiscs due to the bending of TM4, and completely collapses in the IF-asym conformation in MSP1D1 nanodiscs (Figure 3L).

For human Pgp in detergent, we observe a structural response to verapamil in the IF-narrow conformation, mainly manifested with a modest bending of TM10 (∼5 Å at G872) between the residues P866 and D886 (Figure 3G, coral), while TM4 still remains straight (Figure 3G, orange). The IF-wide conformation retains straight helices in both TM4 and TM10 after verapamil binding and could account for an earlier state in substrate recognition, as also recently postulated ^9^. In contrast, the IF-wide conformation of mouse P-gp displays a pronounced bending (∼12 Å at S876) in the equivalent segment of TM10 (P862-D882), under the same conditions (Figure 3J).

The most pronounced effects are seen in human Pgp in MSP2N2 nanodiscs upon addition of verapamil. Here, TM4 displays slightly higher bending (∼12 Å at K234) than in the apo state (Figure 3H, orange helices), while TM10 also bends around residue S880 by ∼12 Å (Figure 3H, coral). In the smaller nanodiscs, the verapamil-bound IF-narrow conformation resembles the conformation resolved in MSP2N2 nanodiscs, with both helices bent at a similar degree (Figure 3I). Similar structures featuring two bent helices were previously resolved in the presence of other substrates and inhibitors ^10,29,33,35^. Previous studies suggested that both TM4 and TM10 bend in response to substrate binding; however, the impact of the hydrophobic environments had not been probed. Our data indicate that TM4 is more influenced by the hydrophobic environment than by the substrate.

### Verapamil binding to human and mouse P-gp

Irrespective of environment and conformation, a single, clear density of verapamil could be resolved in the central binding pocket of the IF-wide and IF-narrow structures of human P-gp, albeit with environment-specific binding modes. An exception is the unusual IF-asym, which did not reveal any verapamil density in the binding site. In the detergent structures, verapamil occupies an almost identical position, with only minor adjustments (Figures 4C and G, Figure S5D and E). In the MSP2N2 (Figure 4D and 4H) and MSP1D1 nanodiscs (Figure 4E and 4I), where TM4 is pre-bent, verapamil also resides in a similar position within the binding pocket (Figure 4K), but with a markedly distinct pose. This is likely caused by the reduced accessible volume within the binding pocket of the nanodisc-reconstituted transporter, as described above (Figure 3L). Similar observations can also be found in the recently published conformations of verapamil-bound P-gp under active turnover conditions ^9^. Regardless of the binding pose, the interaction occurs through similar hydrophobic residues in the drug binding pocket. The primary contacts lie in TM1 (M68 and M69), TM5 (F303 and Y310), TM6 (F336, L339, I340 and F343) and TM12 (F983, M986, and Q990) (Figure 4G, H, I), suggesting that verapamil occupies the M-site, one of three main drug-binding sites (H – site, R - site and M or the modulator site), proposed for P-gp ^32,36^.

**Figure 4.**
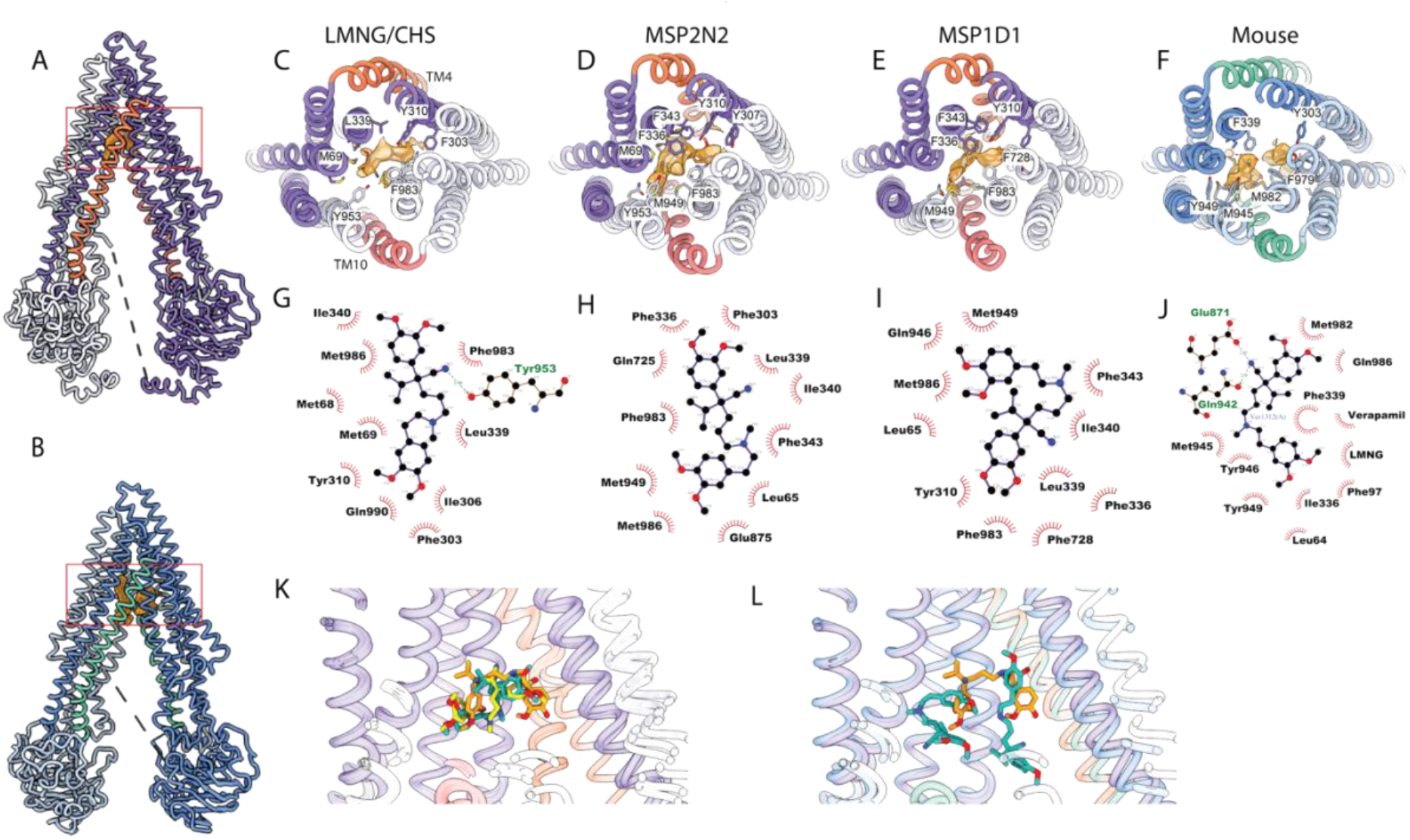
Verapamil binding to human and mouse P-gp. An atomic model of verapamil bound human P-gp in LMNG/CHS in IF-wide conformation. (A) and mouse P-gp in detergent micelles (B). The cryo-EM densities of verapamil in the binding pocket are highlighted in orange. Top views of human P-gp in LMNG/CHS micelles (C, IF-wide), MSP2N2 (D, IF-narrow), MSP1D1 (E, IF-narrow), and mouse P-gp in detergent micelles (F, IF-wide), highlighting one (human) or two molecules (mouse) of verapamil bound in the central binding pocket and the interacting residues. 2D ligand interaction diagram ^37^ of verapamil and the surrounding residues for human P-gp in detergent micelles (G), MSP2N2 (H), MSP1D1 (I), and mouse P-gp in detergent micelles (J). K) Verapamil occupies a similar position in human Pgp, but adapts different poses in different hydrophobic environments, LMNG/CHS (orange), MSP2N2 (yellow), and MSP1D1 (light blue). L) Overlay of the positions and poses of verapamil in human (orange) and mouse P-gp (light blue).

Interestingly, in mouse P-gp, two verapamil molecules were resolved within the substrate-binding pocket in detergent micelles (Figure 4B and F). In addition, an LMNG molecule was identified in the binding pocket, which may have interfered with the verapamil-binding configuration (Figure S5B). Nevertheless, verapamil interacts with similar hydrophobic residues as in human P-gp, including M67, M68, and L64 in TM1; F332, I336, F339, and Q343 in TM6; Q942, M945, and Y949 in TM11; and F974, F979, V978, and M982 in TM12 (Figure 4J and L). Here, one molecule of verapamil is located at the bottom of the M-site, while the second molecule is sitting in both the M and R-sites. Altogether, this suggests that verapamil does not exhibit a single specific binding configuration but rather can adopt different poses, depending on the conformation of the transporter, which could provide a plausible explanation for the polyspecificity and malleability of P-gp.

## Discussion

Transport substrates and the hydrophobic environment influence the ATPase activity of P-gp; however, the molecular mechanisms behind this remain obscure. It has been recognized that TM4 and TM10 are typically bent in substrate-or inhibitor-bound conformations ^8,10,29^, and together with substrate binding decreases the NBD distance, which serves as the main explanation of the stimulated ATPase activity ^38,39^. Additionally, not only substrates and inhibitors, but also the Fab fragment MRK16, induce bending of the TM4 and 10 helices and stimulate ATP hydrolysis ^10^. However, ATPase activity is not only stimulated by substrates but also by the hydrophobic environment ^14^, and the environment itself also impacts the conformational spectrum and NBD separation ^17^. Here, through comparison of detergent-and nanodisc-stabilized P-gp structures in the apo and verapamil-bound states, we reveal distinct roles of TM4 and TM10 in sensing the environment and the substrate and highlight their combined effects on the conformational spectrum and the plasticity within the binding pocket of P-gp.

Our data show that helical bending is not directly correlated with NBD separation. Both IF-wide and IF-narrow conformations are present with bent and straight helices; only the IF-wide+ conformation showed exclusively straight helices. As such, bending of TM4 and TM10 is not required for the spatial approximation of the NBDs, but the likelihood of bent helices in narrower conformations is significantly higher.

While both IF-wide and IF-narrow conformations were detected in the presence of verapamil, narrower conformations were more populated compared to the apo spectrum. This behavior is similar when detergent and nanodisc samples are compared, as previously reported for MsbA ^17^. Consequently, the most pronounced shift in equilibrium towards narrower conformations is observed in the nanodisc-stabilized samples in the presence of verapamil. We observe a good correlation between the bending of TM4 and TM10 and the ATPase activity of the transporter. The low basal ATP hydrolysis in detergent-stabilized apo P-gp coincides with straight TM4 and TM10. The binding of verapamil significantly stimulates ATPase activity, which is associated with a strong reaction of TM10, as evident from the verapamil-bound P-gp structures in detergent. Interestingly, for verapamil-bound mouse P-gp, TM10 bending is much more pronounced compared to the modest bending in human P-gp, which could explain the two-fold higher ATPase activity of mouse P-gp compared to its human counterpart in detergent micelles.

Importantly, in detergent, TM4 remains straight even after verapamil binding, which highlights distinct roles of TM10 and TM4. In the literature, TM4 was observed to be bent in detergent when crystallized with an inhibitor ^40^ and when crosslinked NBDs and an inhibitory UIC2 Fab fragment were used ^33^. However, these observations are likely related to sample preparation. In the first case, the prolonged interaction with the inhibitor during crystal formation likely affected helical conformation. While in the second one, the artificial narrowing of the IF conformation induced by NBD crosslinking or the Fab fragment likely increased the probability of a structural response of TM4.

Across all our data, TM4 bending occurs only in nanodiscs, independent of the drug, and is directly linked to higher basal ATPase activity, as evident from the analysis of nanodisc-embedded human apo P-gp. We concluded that TM4 responds more strongly to environmental changes than to substrate binding. Bending of TM10 is only marginally influenced by the environment, as observed in all apo structures in different nanodiscs. The highest activity is achieved when both helices are bent, as suggested by our and previous data ^9,10^. This requires the presence of the substrate and a specific hydrophobic environment. In our data, the fastest ATPase hydrolysis was measured in MSP2N2 nanodiscs in the presence of verapamil, where the majority of P-gp particles exhibited bending of both TM4 and TM10 helices. Together, our data support a model in which the additive effects that lipid environment and substrate binding exert on TM4 and TM10, respectively, provide a plausible explanation for the changes in ATPase activities, and highlight both helices as an integral part of communication between the TMDs and the NBDs (Figure 5).

**Figure 5.**
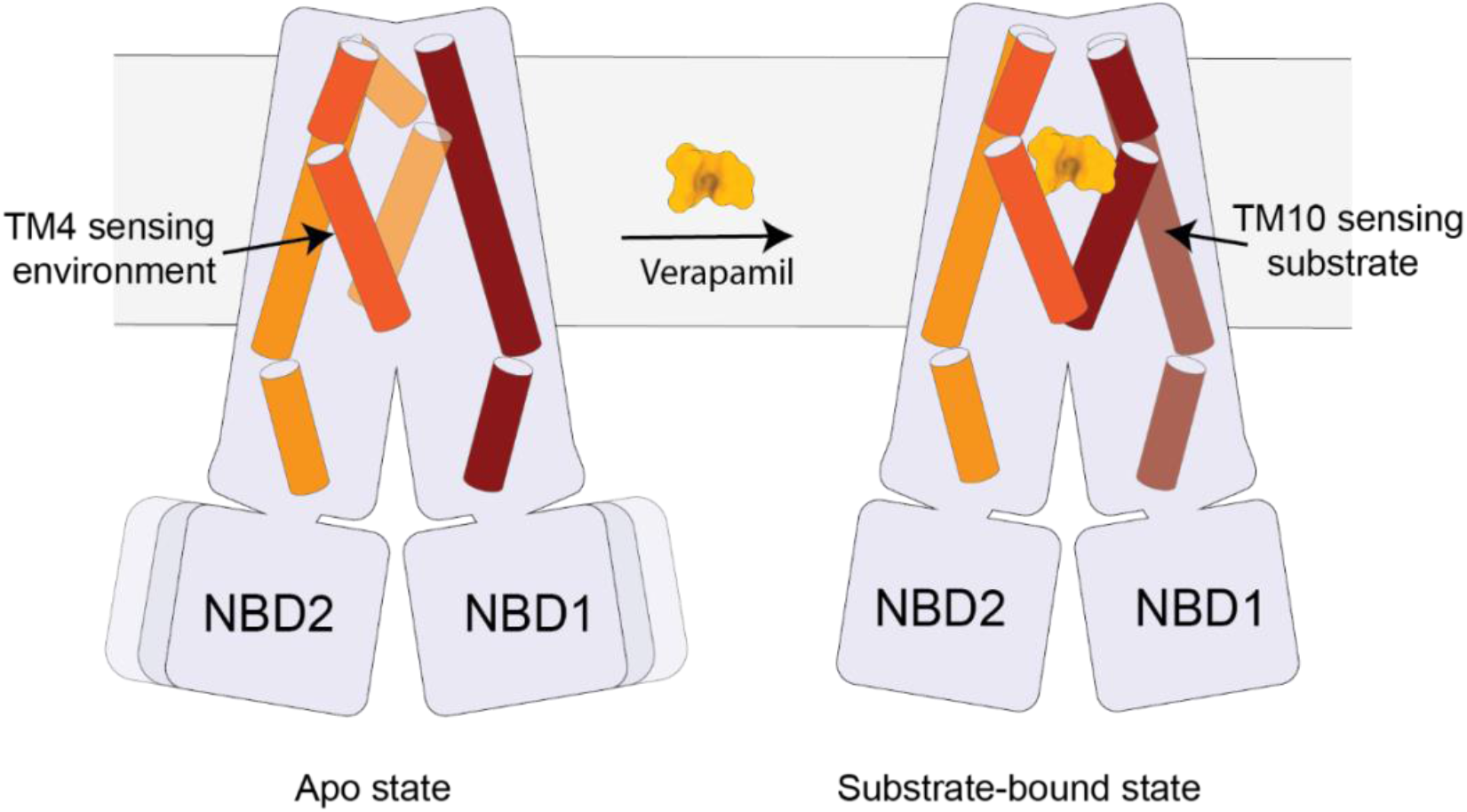
Model describing the dual sensing of TM4 and TM10 to the membrane environment and substrate binding. In the apo state, TM4 exhibits pronounced flexibility and samples different bending configurations in response to the membrane environment. In contrast, TM10 bends once a substrate is present.

The recently reported IF-asym conformation ^8^ aligns well with the sensory functions of TM4 and TM10. In a tight environment characterized by low fluidity and high lipid packing, such as in saposin nanoparticles or small nanodiscs, the IF-asym predominates in the apo configuration. The pronounced kinking of TM4 in this conformation hinders verapamil binding, resulting in a weak response to verapamil stimulation in ATPase assays, as more than half of the particles remain in this conformation in the presence of verapamil. Importantly, the addition of a substrate can rescue the typical IF conformation, as evidenced by the increase in the fraction of the IF-narrow conformation from 14 % in the apo-state to 48 % once verapamil is added.

Furthermore, we noticed a preferred orientation of P-gp within our nanodisc preparation that correlates with the appearance of either IF-asym or typical IF conformations. When TM4 faces the center of the nanodisc, IF-asym conformers are observed; when TM4 faces the periphery of the disc, the typical IF conformation is obtained. We suggest that this observation can be explained by different regions of lipid populations that have been reported for nanodiscs: the center of the nanodisc is composed of more tightly packed lipids, whereas disorder increases towards the periphery. Moreover, the thickness and membrane stiffness are the highest in the middle of the small nanodisc ^41,42^. Consequently, the high-order and membrane stiffness at the center of the small nanodisc may stabilize TM4 in its unusual IF-asym conformation. While the soft environment at the nanodisc edge may allow TM4 to be more flexible. Intriguingly, both stiffness and membrane thickness decrease at the center of the larger nanodiscs, MSP2N2, which explains the absence of the IF-asym conformation in this environment ^42^.

The observed differences in verapamil’s binding poses between detergent and nanodiscs likely arise from conformational variations already present in the apo states across the respective environments. TM4 has the most profound impact on reshaping the binding pocket and, together with TM10, reduces the accessible volume in the nanodisc. Therefore, verapamil likely alters its pose to accommodate the plastic binding site. This suggests that the substrate does not exhibit a specific binding configuration but rather adopts a different pose depending on the transporter’s conformation, which in turn is influenced by the membrane environment.

Utilizing different hydrophobic environments allowed us to decipher distinct roles for TM4 and TM10 of P-gp. The pronounced response of TM4 to the membrane mimetic suggests a primary role in sensing the surrounding environment; in contrast, TM10 primarily senses substrate binding. This dual regulation of P-gp impacts its conformational spectrum and provides a new view on its functional plasticity.

## METHODS

### Expression and purification of P-gp

Human wild-type MDR1 cDNA (GenBank: BC130424.1) containing the S400N polymorphism and the sequence AAAAAA immediately upstream of the ATG initiation codon was excised from the pHIL-hMDR1.4 vector ^43^ and cloned into the pPICZ vector (Invitrogen/ThermoFisher) using In-Fusion seamless cloning (Takarabio.com). Sequences encoding TEV-cleavable Twin-Strep and His_6_ purification tags were added immediately after the last amino acid as follows: ctc gag gaa aac ctg tat ttt cag ggc gga ggc gcc tct ggt ggt tct tgg tct cac cca caa ttc gag aag gcg gcc gcc ggc ggt ggt tct ggc ggt ggt tct tgg tct cac cca caa ttc gag aag ggt tct ggt cac cac cat cac cac cat tga. This engineered tag is identical to the C-terminus of mouse Mdr1a P-gp (accession number NM_011076, GenBank JF83415) previously published ^14,44^. P-gp was expressed in *Pichia pastoris*, and the proteins were purified from microsomal membranes following established protocols ^14,44,45^. Briefly, P-gp microsomal membranes were solubilized in 1% DDM/0.1% CHS and purified by tandem affinity chromatography on Ni-NTA (Qiagen) and Strep-Tactin XT resins (IBA Lifesciences) in buffers containing 0.02% LMNG/0.004% CHS. The eluted fractions were further purified using size-exclusion chromatography (SEC) with either Superdex 200 Increase 10/300 (Cytiva) or Superose 6 Increase 10/300 (Cytiva) (Figure S1A, C) in SEC buffer (200 mM NaCl, 20 mM Tris/HCl, pH 7.4, 1 mM DTT) containing 0.02% LMNG/0.004% CHS. Peak fractions were pooled together and concentrated for further experiments.

For nanodisc reconstitution, P-gp was purified in buffers containing 0.05% DDM and 0.005% CHS. Briefly, solubilized membranes were loaded onto a Histrap™ HP (Cytiva) column, which was subsequently washed with 20 column volumes of wash buffer (500 mM NaCl, 50 mM Tris/HCl, pH 8.0, 10% glycerol, 50 mM imidazole, 1 mM DTT, and 0.05% DDM/0.005% CHS). The protein was then eluted with 300 mM imidazole in the same buffer. Eluted fractions were pooled, concentrated, and loaded onto a Superdex 200 increase 10/300 or Superose 6 increase 10/300 column in SEC buffer containing 0.05% DDM/0.005% CHS for further polishing (Figure S1B). Peak fractions were pooled and concentrated for reconstitution into nanodiscs.

### Reconstitution of P-gp in nanodiscs

Brain Polar Lipid Extract (Avanti Research, A84101) and cholesterol were mixed at a ratio of 80:20 (w/w) from a chloroform stock (25 mg/ml). Chloroform was evaporated in a rotary evaporator (Heidolph Hei Vap). The lipid film was resuspended in 150 mM NaCl and 20 mM Tris/HCl (pH 7.4) to form liposomes at a final concentration of 4 mg/ml. Liposomes were dissolved using 1% Triton X-100 and mixed with the purified P-gp and the MSP protein to achieve a molar ratio (P-gp: MSP: lipid) of 1:10:1000 or 1:10:350 for MSP2N2 or MSP1D1, respectively. The mixture was incubated with approximately 200 mg of Biobeads SM2 at 4°C overnight in a roller shaker. The nanodiscs were centrifuged at 14,000 × g in a tabletop centrifuge for 30 minutes at 4 °C to remove any aggregates or proteoliposomes. To separate P-gp loaded nanodiscs from empty nanodiscs, the supernatant was loaded onto Ni-NTA beads that bind the His-tagged P-gp. The beads were washed with (20 mM Tris/HCl, pH 7.5, 200 mM NaCl, 40 mM imidazole), and P-gp nanodiscs were eluted with the same buffer containing 300 mM imidazole. The elution fractions were concentrated and loaded onto Superose 6 increase 10/300 in the case of MSP2N2 or Superdex 200 increase 10/300 in the case of MSP1D1 nanodiscs (Figures S1D, E). The peak fractions were collected and concentrated to approximately 2 mg/ml for cryo-EM studies.

### ATPase activity assay

The ATPase activity of P-gp was measured using a molybdate-based colorimetric assay as previously described ^46^, with minor adjustments. Briefly, 2 µg of LMNG/CHS-purified P-gp or 1 µg of nanodisc-reconstituted P-gp was incubated with ATP cocktail (2 mM ATP, 4 mM MgCl_2_, in 50 mM Tris-HCl, pH 7.4 buffer) in the absence (for basal activity) or presence of verapamil (15–1,000 µM) at 37°C for 30 min. Verapamil-HCl stocks (100 mM) were prepared in 20 mM Hepes /KOH, pH 6.8. Reactions were stopped by diluting the samples 1:1 with 12% SDS (w/v). 12% (w/v) ascorbic acid and 2% (w/v) ammonium molybdate in 1 M HCl were added, and the mixture was incubated for 5 min at room temperature. Subsequently, 25 mM sodium citrate and 2% (w/v) sodium meta-arsenite in 2% (w/v) acetic acid were added, and samples were incubated for an additional 2 min. Absorbance was measured at 850 nm using Infinite^R^ M Nano + microplate reader (TECAN). Phosphate concentrations were calculated from a standard curve (0-60 nmol). Data are shown as mean ± SEM (n = 3).

### Cryo-EM sample preparation

For cryo-EM experiments, 5 mg/ml of LMNG/CHS-purified P-gp was used for detergent-based preparations, or 2 mg/ml of nanodiscs-reconstituted P-gp. For verapamil-bound samples, human P-gp was incubated with 250 µM verapamil, while mouse P-gp was incubated with 50 µM verapamil for 30 min at room temperature. C-flat CF-1.2/1.3 Cu 300 mesh grids (Protochips) were glow-discharged using a PELCO easiGlow (15 mA, 120 s). A sample volume of 3 µl was applied to grids and vitrified using a Vitrobot Mark IV (Thermo Fisher) at 4°C and 100% relative humidity. Cryo-EM data were acquired on a 200-kV Glacios microscope (Thermo Fisher) equipped with a Selectris energy filter set to a slit width of 10-eV. Movies were recorded using a Falcon 4i direct electron detector in EER format at a nominal magnification of 165,000, corresponding to a physical pixel size of 0.68 Å, with a total dose of 50 e^−^/Å^2^. All datasets were collected automatically using EPU v2.9 with a defocus range of −0.6 to −1.8 µm.

### Cryo-EM data processing

All datasets were processed using cryoSPARC (v.4.2-4). A representative processing workflow is shown in Figures S8 and S9. Movies were preprocessed with patch-based motion correction and patch-based CTF estimation and filtered by the CTF fit estimates using a cutoff at 5 Å in cryoSPARC live. Details about the number of collected images and picked particles, the number of particles in the final map, and the resolutions are listed in Table. S1.

All datasets were processed according to the workflow outlined below. Particle positions from blob-picker, template picker, and particles picked with Topaz were combined, filtered for removal of duplicate particles, extracted with 4x binning, and subjected to two rounds of 2D classification. Well-defined 2D classes were selected and used for multi-class ab initio reconstruction. Multiple rounds of ab initio reconstruction, heterogeneous refinement (HR) and non-uniform refinement (NUR) were used to generate a substack of the best particles. Re-extracted particles without binning were used for a final NUR or further local refinement. Reported B-factors result from unsupervised auto-sharpening during refinement in cryoSPARC.

### Model building, refinement, and validation

The initial atomic models were generated with AlphaFold ^47^, and were fitted into the densities as rigid bodies in ChimeraX ^48^. The structure was manually refined using Coot (v.0.9) ^49^ and iteratively refined using phenix.real_space_refine within Phenix (v.1.19) ^50^. Validation reports were automatically generated using MolProbity ^51^ within Phenix.

## Date availability

The cryo-EM density maps supporting this study have been deposited in the Electron Microscopy Data Bank under the accession numbers EMD-56584, EMD-56585, EMD-56586, EMD-56587, EMD-56588, EMD-56589, EMD-56590, EMD-56591, EMD-56592, EMD-56593, EMD-56594, EMD-56595, and EMD-56596. The atomic coordinates of the corresponding models have been deposited in the Protein Data Bank with the accession numbers 28KU, 28KV, 28KX, 28KY, 28KZ, 28LA, 28LB, 28LC, 28LD, 28LE, 28LF, and 28LG.

## Acknowledgements

This work was supported by a grant of the DFG 37802001, the SFB 1557 - 467522186, the DFG-FUGG 455249646, and the RTG 2900 - 501879556 (all to AM) and the National Institute of General Medical Sciences R01 GM141216 (ILU).

## Author Contribution

MK conceived and designed all experiments together with DJ, IU, and AM. MK, JHS, and VJM conducted the cryo-EM analyses, model building, and validation with support from KS, KP, and DJ. MK, VJM, NT, and DM performed biochemical analyses with support from IU. MK, DJ, DS, IU, and AM interpreted the data. The manuscript was written by MK, DJ, IU, and AM with contributions from all the authors. Funding acquisition was done by IU and AM.

## References

1. Seelig, A. (2020). P-Glycoprotein: One Mechanism, Many Tasks and the Consequences for Pharmacotherapy of Cancers. Front. Oncol. 10, 576559. 10.3389/FONC.2020.576559.

2. Chufan, E.E., Sim, H.-M., and Ambudkar, S. V (2015). Molecular Basis of the Polyspecificity of P-Glycoprotein (ABCB1). In ABC Transporters and Cancer Advances in cancer research. (Elsevier), pp. 71–96.

3. Thomas, C., Aller, S.G., Beis, K., Carpenter, E.P., Chang, G., Chen, L., Dassa, E., Dean, M., Duong Van Hoa, F., Ekiert, D., et al. (2020). Structural and functional diversity calls for a new classification of ABC transporters. FEBS Lett. 594, 3767–3775. 10.1002/1873-3468.13935.

4. Alam, A., and Locher, K.P. (2023). Structure and Mechanism of Human ABC Transporters. Annu. Rev. Biophys. 52, 275–300. 10.1146/ANNUREV-BIOPHYS-111622-091232/CITE/REFWORKS.

5. Jardetzky, O. (1966). Simple allosteric model for membrane pumps. Nature 211, 969–970.

6. Hofmann, S., Januliene, D., Mehdipour, A.R., Thomas, C., Stefan, E., Brüchert, S., Kuhn, B.T., Geertsma, E.R., Hummer, G., Tampé, R., et al. (2019). Conformation space of a heterodimeric ABC exporter under turnover conditions. Nature 2019 571:7766 571, 580–583. 10.1038/s41586-019-1391-0.

7. Ambudkar, S. V. (1998). Drug-stimulatable ATPase activity in crude membranes of human MDR1-transfected mammalian cells. Methods Enzymol. 292, 504–514. 10.1016/S0076-6879(98)92039-0.

8. Kurre, D., Dang, P.X., Le, L.T.M., Gadkari, V. V., and Alam, A. (2025). Structural insights into binding-site access and ligand recognition by human ABCB1. EMBO Journal 44, 991–1006. 10.1038/s44318-025-00361-z.

9. Culbertson, A.T., and Liao, M. (2025). Cryo-EM of human P-glycoprotein reveals an intermediate occluded conformation during active drug transport. Nature Communications 16. 10.1038/s41467-025-58561-4.

10. Nosol, K., Romane, K., Irobalieva, R.N., Alam, A., Kowal, J., Fujita, N., and Locher, K.P. (2020). Cryo-EM structures reveal distinct mechanisms of inhibition of the human multidrug transporter ABCB1. Proc. Natl. Acad. Sci. U. S. A. 117, 26245–26253. 10.1073/PNAS.2010264117;WGROUP:STRING:PUBLICATION.

11. Moeller, A., Lee, S.C., Tao, H., Speir, J.A., Chang, G., Urbatsch, I.L., Potter, C.S., Carragher, B., and Zhang, Q. (2015). Distinct Conformational Spectrum of Homologous Multidrug ABC Transporters. Structure 23, 450–460. 10.1016/J.STR.2014.12.013.

12. Gewering, T., Waghray, D., Parey, K., Jung, H., Tran, N.N., Zapata, J., Zhao, P., Chen, H., Januliene, D., Hummer, G., et al. (2024). Tracing the substrate translocation mechanism in P-glycoprotein. Elife 12. 10.7554/elife.90174.

13. Shukla, S., Abel, B., Chufan, E.E., and Ambudkar, S. V. (2017). Effects of a detergent micelle environment on P-glycoprotein (ABCB1)-ligand interactions. Journal of Biological Chemistry 292, 7066–7076. 10.1074/JBC.M116.771634.

14. Tran, N.N.B., Bui, A.T.A., Jaramillo-Martinez, V., Weber, J., Zhang, Q., and Urbatsch, I.L. (2023). Lipid environment determines the drug-stimulated ATPase activity of P-glycoprotein. Front. Mol. Biosci. 10, 1141081. 10.3389/FMOLB.2023.1141081/BIBTEX.

15. Song, G., Zhang, S., Tian, M., Zhang, L., Guo, R., Zhuo, W., and Yang, M. (2021). Molecular insights into the human ABCB6 transporter. Cell Discovery 2021 7:1 7, 55-. 10.1038/s41421-021-00284-z.

16. Plummer-Medeiros, A.M., Culbertson, A.T., Morales-Perez, C.L., and Liao, M. (2023). Activity and Structural Dynamics of Human ABCA1 in a Lipid Membrane. J. Mol. Biol. 435, 168038. 10.1016/J.JMB.2023.168038.

17. Hoffmann, L., Baier, A., Jorde, L., Kamel, M., Schäfer, J.H., Schnelle, K., Scholz, A., Shvarev, D., Wong, J.E.M.M., Parey, K., et al. (2025). The ABC transporter MsbA in a dozen environments. Structure 33, 916-923.e4. 10.1016/J.STR.2025.02.002.

18. Galazzo, L., Meier, G., Januliene, D., Parey, K., De Vecchis, D., Striednig, B., Hilbi, H., Schäfer, L. V., Kuprov, I., Moeller, A., et al. (2022). The ABC transporter MsbA adopts the wide inward-open conformation in E. coli cells. Sci. Adv. 8, 6845. 10.1126/SCIADV.ABN6845.

19. Takeuchi, T., Yoshitomi, S., Higuchi, T., Ikemoto, K., Niwa, S.I., Ebihara, T., Katoh, M., Yokoi, T., and Asahi, S. (2006). Establishment and Characterization of the Transformants Stably-Expressing MDR1 Derived from Various Animal Species in LLC-PK1. Pharmaceutical Research 2006 23:7 23, 1460–1472. 10.1007/S11095-006-0285-7.

20. Baltes, S., Gastens, A.M., Fedrowitz, M., Potschka, H., Kaever, V., and Löscher, W. (2007). Differences in the transport of the antiepileptic drugs phenytoin, levetiracetam and carbamazepine by human and mouse P-glycoprotein. Neuropharmacology 52, 333–346. 10.1016/J.NEUROPHARM.2006.07.038.

21. Yamazaki, M., Neway, W.E., Ohe, T., Chen, I.W., Rowe, J.F., Hochman, J.H., Chiba, M., and Lin, J.H. (2001). In Vitro Substrate Identification Studies for P-glycoprotein-Mediated Transport: Species Difference and Predictability of in Vivo Results. J. Pharmacol. Exp. Ther. 296, 723–735. 10.1016/S0022-3565(24)38809-3.

22. Nandigama, K., Lusvarghi, S., Shukla, S., and Ambudkar, S. V. (2019). Large-scale purification of functional human P-glycoprotein (ABCB1). Protein Expr. Purif. 159, 60–68. 10.1016/J.PEP.2019.03.002.

23. Clay, A.T., Lu, P., and Sharom, F.J. (2015). Interaction of the P-Glycoprotein Multidrug Transporter with Sterols. Biochemistry 54, 6586–6597. 10.1021/ACS.BIOCHEM.5B00904.

24. Kimura, Y., Kodan, A., Matsuo, M., and Ueda, K. (2007). Cholesterol fill-in model: mechanism for substrate recognition by ABC proteins. Journal of Bioenergetics and Biomembranes 2007 39:5 39, 447–452. 10.1007/S10863-007-9109-7.

25. Kimura, Y., Kioka, N., Kato, H., Matsuo, M., and Ueda, K. (2007). Modulation of drug-stimulated ATPase activity of human MDR1/P-glycoprotein by cholesterol. Biochemical Journal 401, 597–605. 10.1042/BJ20060632.

26. Doige, C.A., Yu, X., and Sharom, F.J. (1993). The effects of lipids and detergents on ATPase-active P-glycoprotein. Biochimica et Biophysica Acta (BBA) - Biomembranes 1146, 65–72. 10.1016/0005-2736(93)90339-2.

27. Li-Blatter, X., Nervi, P., and Seelig, A. (2009). Detergents as intrinsic P-glycoprotein substrates and inhibitors. Biochim. Biophys. Acta Biomembr. 1788, 2335–2344. 10.1016/j.bbamem.2009.07.010.

28. Seelig, A., and Li-Blatter, X. (2023). P-glycoprotein (ABCB1) - weak dipolar interactions provide the key to understanding allocrite recognition, binding, and transport. Cancer Drug Resistance 6, 1. 10.20517/CDR.2022.59.

29. Alam, A., Kowal, J., Broude, E., Roninson, I., and Locher, K.P. (2019). Structural insight into substrate and inhibitor discrimination by human P-glycoprotein. Science (1979). 363, 753–756. 10.1126/SCIENCE.AAV7102.

30. Singh, B.N., Ellrodt, G., and Peter, C.T. (1978). Verapamil. Drugs 15, 169–197.

31. Tsuruo, T., Iida, H., Tsukagoshi, S., and research, Y.S. (1981). Overcoming of Vincristine Resistance in P388 Leukemia in Vivo and in Vitro through Enhanced Cytotoxicity of Vincristine and Vinblastine by Verapamil. Cancer Res.

32. Ferreira, R.J., Ferreira, M.J.U., and Dos Santos, D.J.V.A. (2013). Molecular Docking Characterizes Substrate-Binding Sites and Efflux Modulation Mechanisms within P-Glycoprotein. J. Chem. Inf. Model. 53, 1747–1760. 10.1021/CI400195V.

33. Alam, A., Küng, R., Kowal, J., McLeod, R.A., Tremp, N., Broude, E. V., Roninson, I.B., Stahlberg, H., and Locher, K.P. (2018). Structure of a zosuquidar and UIC2-bound human-mouse chimeric ABCB1. Proc. Natl. Acad. Sci. U. S. A. 115, E1973–E1982. 10.1073/pnas.1717044115.

34. Urgaonkar, S., Nosol, K., Said, A.M., Nasief, N.N., Bu, Y., Locher, K.P., Lau, J.Y.N., and Smolinski, M.P. (2022). Discovery and Characterization of Potent Dual P-Glycoprotein and CYP3A4 Inhibitors: Design, Synthesis, Cryo-EM Analysis, and Biological Evaluations. J. Med. Chem. 65, 191–216. 10.1021/ACS.JMEDCHEM.1C01272.

35. Hamaguchi-Suzuki, N., Adachi, N., Moriya, T., Yasuda, S., Kawasaki, M., Suzuki, K., Ogasawara, S., Anzai, N., Senda, T., and Murata, T. (2024). Cryo-EM structure of P-glycoprotein bound to triple elacridar inhibitor molecules. Biochem. Biophys. Res. Commun. 709, 149855. 10.1016/J.BBRC.2024.149855.

36. Aller, S.G., Yu, J., Ward, A., Weng, Y., Chittaboina, S., Zhuo, R., Harrell, P.M., Trinh, Y.T., Zhang, Q., Urbatsch, I.L., et al. (2009). Structure of P-glycoprotein reveals a molecular basis for poly-specific drug binding. Science (1979). 323, 1718–1722. 10.1126/SCIENCE.1168750.

37. Laskowski, R.A., and Swindells, M.B. (2011). LigPlot+: Multiple Ligand–Protein Interaction Diagrams for Drug Discovery. J. Chem. Inf. Model. 51, 2778–2786. 10.1021/CI200227U.

38. Loo, T.W., Bartlett, M.C., and Clarke, D.M. (2003). Drug Binding in Human P-glycoprotein Causes Conformational Changes in Both Nucleotide-binding Domains. Journal of Biological Chemistry 278, 1575–1578. 10.1074/jbc.M211307200.

39. Zoghbi, M.E., Mok, L., Swartz, D.J., Singh, A., Fendley, G.A., Urbatsch, I.L., and Altenberg, G.A. (2017). Substrate-induced conformational changes in the nucleotide-binding domains of lipid bilayer-associated P-glycoprotein during ATP hydrolysis. Journal of Biological Chemistry 292, 20412–20424. 10.1074/jbc.M117.814186.

40. Szewczyk, P., Tao, H., McGrath, A.P., Villaluz, M., Rees, S.D., Lee, S.C., Doshi, R., Urbatsch, I.L., Zhang, Q., and Chang, G. (2015). Snapshots of ligand entry, malleable binding and induced helical movement in P-glycoprotein. Acta Crystallogr. D Biol. Crystallogr. 71, 732–741. 10.1107/S1399004715000978.

41. Stepien, P., Augustyn, B., Poojari, C., Galan, W., Polit, A., Vattulainen, I., Wisnieska-Becker, A., and Rog, T. (2020). Complexity of seemingly simple lipid nanodiscs. Biochimica et Biophysica Acta (BBA) - Biomembranes 1862, 183420. 10.1016/J.BBAMEM.2020.183420.

42. Schachter, I., Allolio, C., Khelashvili, G., and Harries, D. (2020). Confinement in Nanodiscs Anisotropically Modifies Lipid Bilayer Elastic Properties. J. Phys. Chem. B 124, 7166–7175. 10.1021/ACS.JPCB.0C03374.

43. Urbatsch, I.L., Gimi, K., Wilke-Mounts, S., Lerner-Marmarosh, N., Rousseau, M.E., Gros, P., and Senior, A.E. (2001). Cysteines 431 and 1074 Are Responsible for Inhibitory Disulfide Cross-linking between the Two Nucleotide-binding Sites in Human P-glycoprotein. Journal of Biological Chemistry 276, 26980–26987. 10.1074/jbc.M010829200.

44. Swartz, D.J., Mok, L., Botta, S.K., Singh, A., Altenberg, G.A., and Urbatsch, I.L. (2014). Directed evolution of P-glycoprotein cysteines reveals site-specific, non-conservative substitutions that preserve multidrug resistance. Biosci. Rep. 34, e00116. 10.1042/BSR20140062.

45. Bai, J., Swartz, D.J., Protasevich, I.I., Brouillette, C.G., Harrell, P.M., Hildebrandt, E., Gasser, B., Mattanovich, D., Ward, A., Chang, G., et al. (2011). A Gene Optimization Strategy that Enhances Production of Fully Functional P-Glycoprotein in Pichia pastoris. PLoS One 6, e22577. 10.1371/JOURNAL.PONE.0022577.

46. Mi, W., Li, Y., Yoon, S.H., Ernst, R.K., Walz, T., and Liao, M. (2017). Structural basis of MsbA-mediated lipopolysaccharide transport. Nature 2017 549:7671 549, 233–237. 10.1038/nature23649.

47. Jumper, J., Evans, R., Pritzel, A., Green, T., Figurnov, M., Ronneberger, O., Tunyasuvunakool, K., Bates, R., Žídek, A., Potapenko, A., et al. (2021). Highly accurate protein structure prediction with AlphaFold. Nature 2021 596:7873 596, 583–589. 10.1038/s41586-021-03819-2.

48. Pettersen, E.F., Goddard, T.D., Huang, C.C., Meng, E.C., Couch, G.S., Croll, T.I., Morris, J.H., and Ferrin, T.E. (2021). UCSF ChimeraX: Structure visualization for researchers, educators, and developers. Protein Science 30, 70–82. 10.1002/PRO.3943.

49. Emsley, P., and Cowtan, K. (2004). Coot: model-building tools for molecular graphics. Acta Crystallogr. D Biol. Crystallogr. 60, 2126–2132. 10.1107/S0907444904019158.

50. Liebschner, D., Afonine, P. V., Baker, M.L., Bunkoczi, G., Chen, V.B., Croll, T.I., Hintze, B., Hung, L.W., Jain, S., McCoy, A.J., et al. (2019). Macromolecular structure determination using X-rays, neutrons and electrons: recent developments in Phenix. Acta Crystallogr. D Struct. Biol. 75, 861–877. 10.1107/S2059798319011471.

51. Chen, V.B., Arendall, W.B., Headd, J.J., Keedy, D.A., Immormino, R.M., Kapral, G.J., Murray, L.W., Richardson, J.S., and Richardson, D.C. (2009). MolProbity: all-atom structure validation for macromolecular crystallography. Acta Crystallogr. D Biol. Crystallogr. 66, 12–21. 10.1107/S0907444909042073.

