## Supplements for "A dual sensor regulates P-glycoprotein’s structural plasticity"

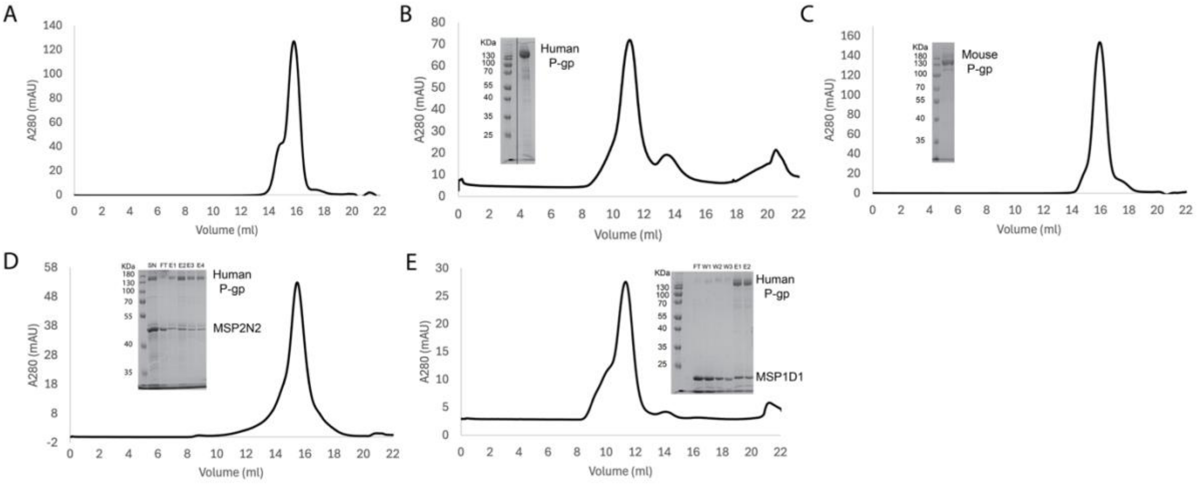

**Figure S1 Human and mouse P-gp purification and nanodisc reconstitution.** Size-exclusion chromatography of A) human P-gp purified in LMNG/CHS and B) DDM/CHS, and of C) mouse P-gp purified in LMNG/CHS. D) Chromatograms and SDS-PAGE analysis of human P-gp reconstitution in MSP2N2 nanodiscs, where SN corresponds to the supernatant after centrifugation of the nanodiscs mixture, FT contains proteins and "empty" nanodiscs not bound to Ni-NTA resin, E1-E4 represent the elution fraction of P-gp-loaded nanodiscs. E) Reconstitution of human P-gp in MSP1D1 nanodiscs, where FT and W1-W3 contain "empty" nanodiscs washed off the resin and E1, E2 represent the elution fractions of P-gp-loaded MSP1D1 nanodiscs.

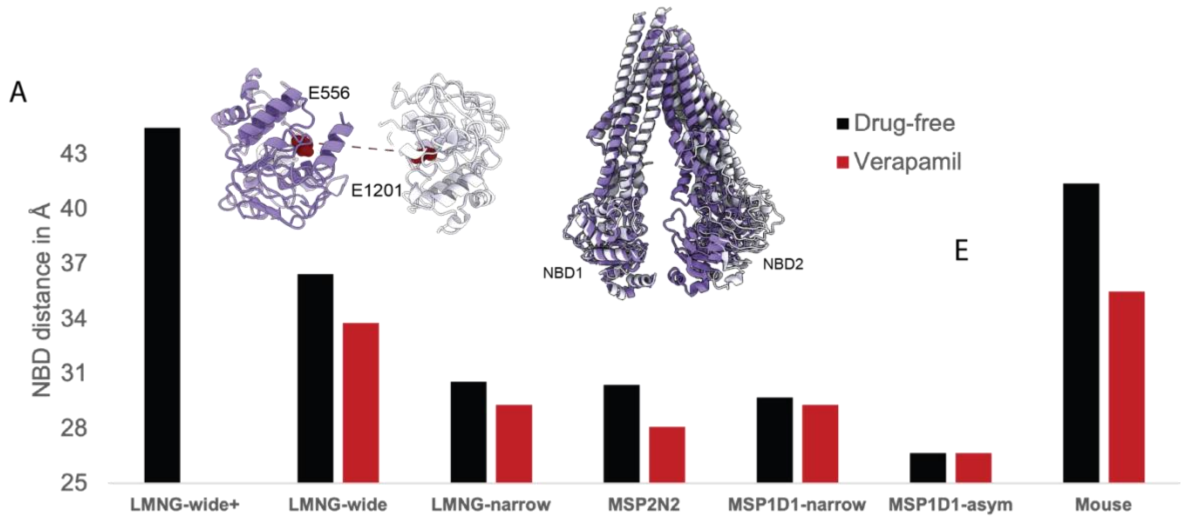

**Figure S2 Inter-NBD distance of P-gp.** The inter-NBD distances in the different conformations of P-gp in detergent micelles and nanodisc as measured between residue E556 and E1201 residues in NBD1 and NBD2 of human P-gp and the equivalent residues in mouse P-gp (E552-E1197).

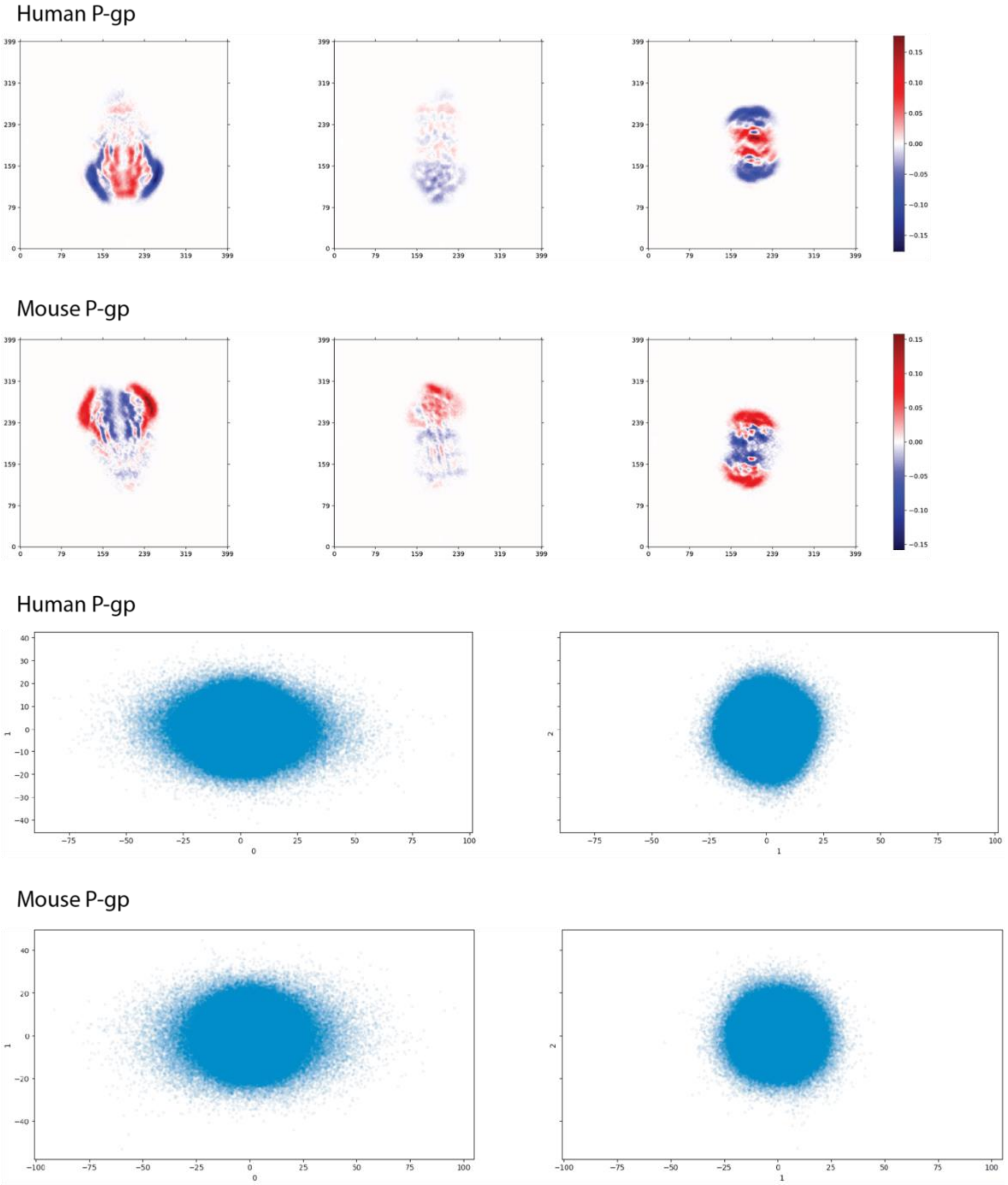

**Figure S3 3D Variability analysis for human and mouse P-gp in the apo state in detergent micelles.** 3D variability analysis shows that both human and mouse P-gp exhibit flexibility, which is more pronounced in human P-gp. The upper panel depicts the motion described by component 0 of 3DVA, displaying the NBDs moving toward each other. Mouse P-gp predominantly resides in wide conformations with a different degree of NBD distances, while human P-gp undergoes movement from wide to narrow conformations. The lower panel shows the reaction coordinate distribution of particles along components 0, 1, and 2 from the

3D variability analysis for both human and mouse P-gp, showing a wider spread of particles along component 0, indicating higher flexibility.

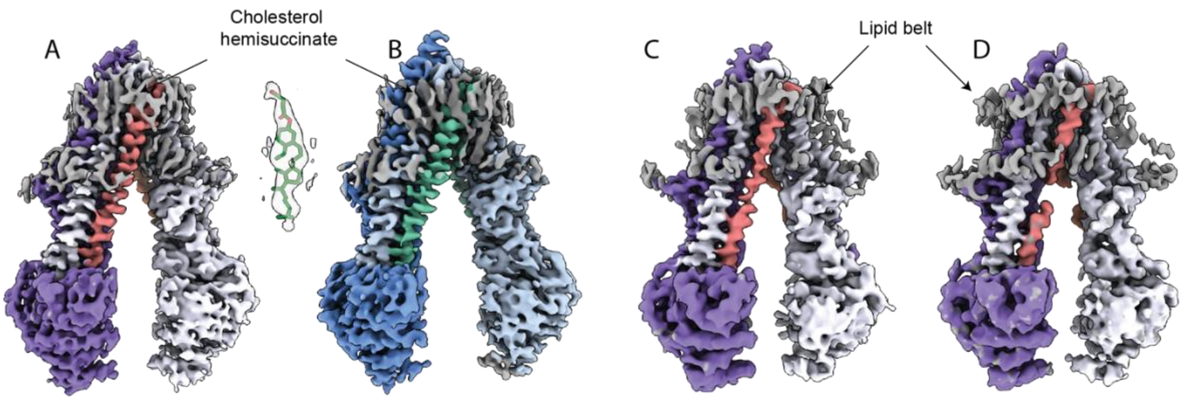

**Figure S4 Lipid and CHS molecules surround P-glycoprotein.** Cholesterol hemisuccinate surrounds the membrane-embedded transmembrane segments of both human (A) and mouse (B) P-gp in detergent micelles. Lipid molecules surround P-glycoprotein in both MSP2N2 (C) and MSP1D1 (D) nanodiscs.

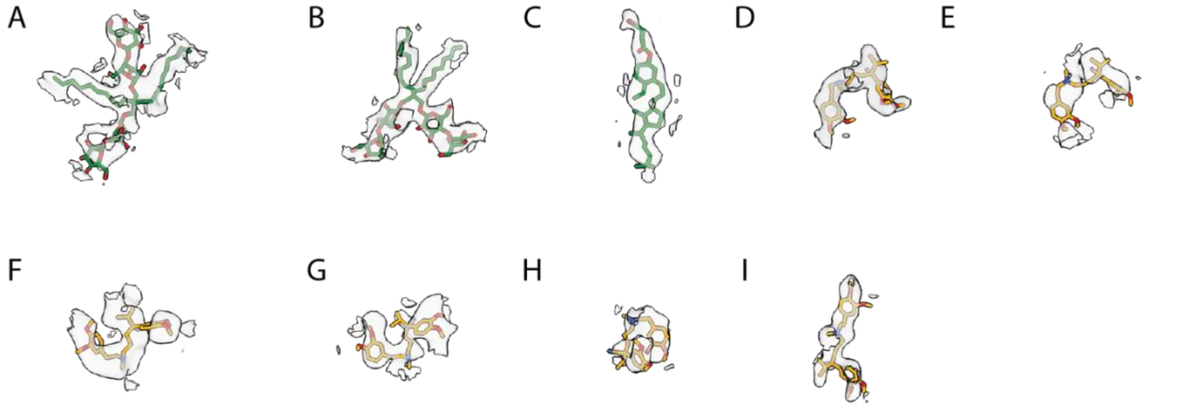

**Figure S5 Cryo-EM density for the resolved substrates and detergent molecules.** A) LMNG molecule identified in the cavity of human P-gp in IF-wide, and B) of verapamil-bound mouse P-gp in detergent micelles. C) An example of a CHS molecule bound to human P-gp in IF-wide conformation. D) verapamil bound to Human P-gp in detergent micelles in the IF-wide conformation. Cryo-EM density of verapamil bound to the IF-narrow conformations of human P-gp in E) detergent micelles, F) in MSP2N2 nanodisc, and G) in MSP1D1 nanodsics. H) and I) EM density of the two molecules of verapamil identified in the binding pocket of mouse P-gp in detergent micelles.

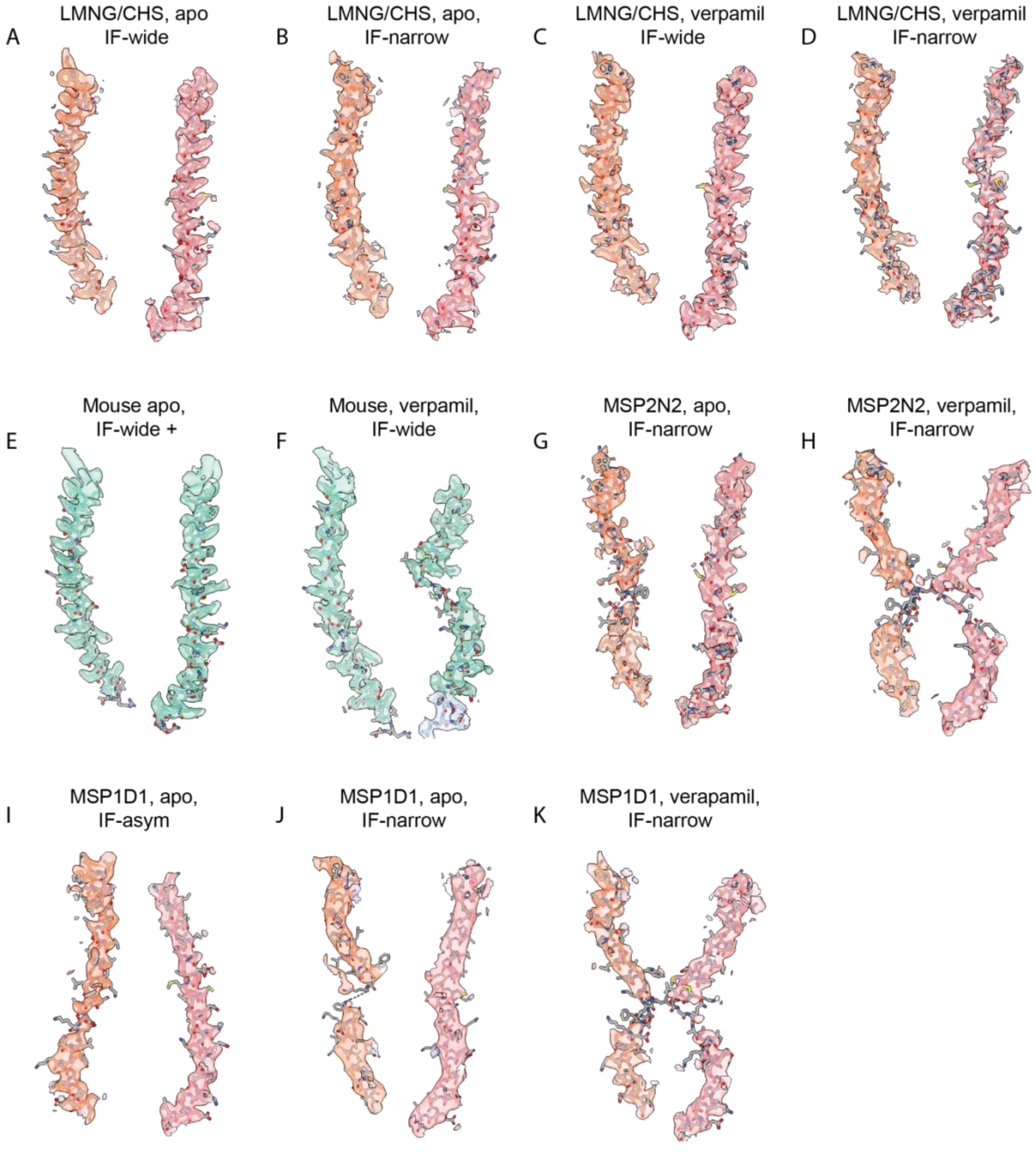

**Figure S6 Local cryo-EM densities of TM4 and TM10.** Human TM4 and TM10 helices are shown in orange and coral, and mouse helices are in green.

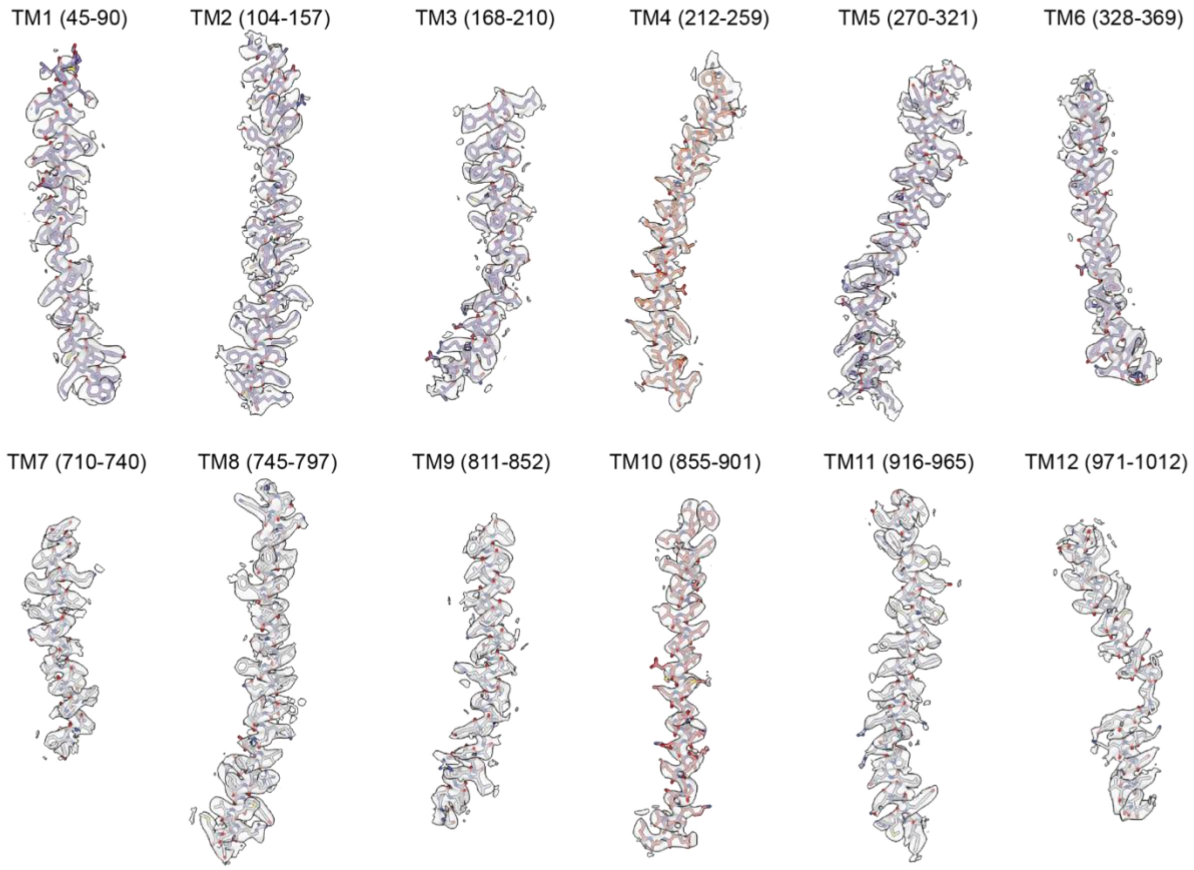

**Figure S7 Local cryo-EM map for all the transmembrane helices of apo human P-gp in the IF-wide conformation.**

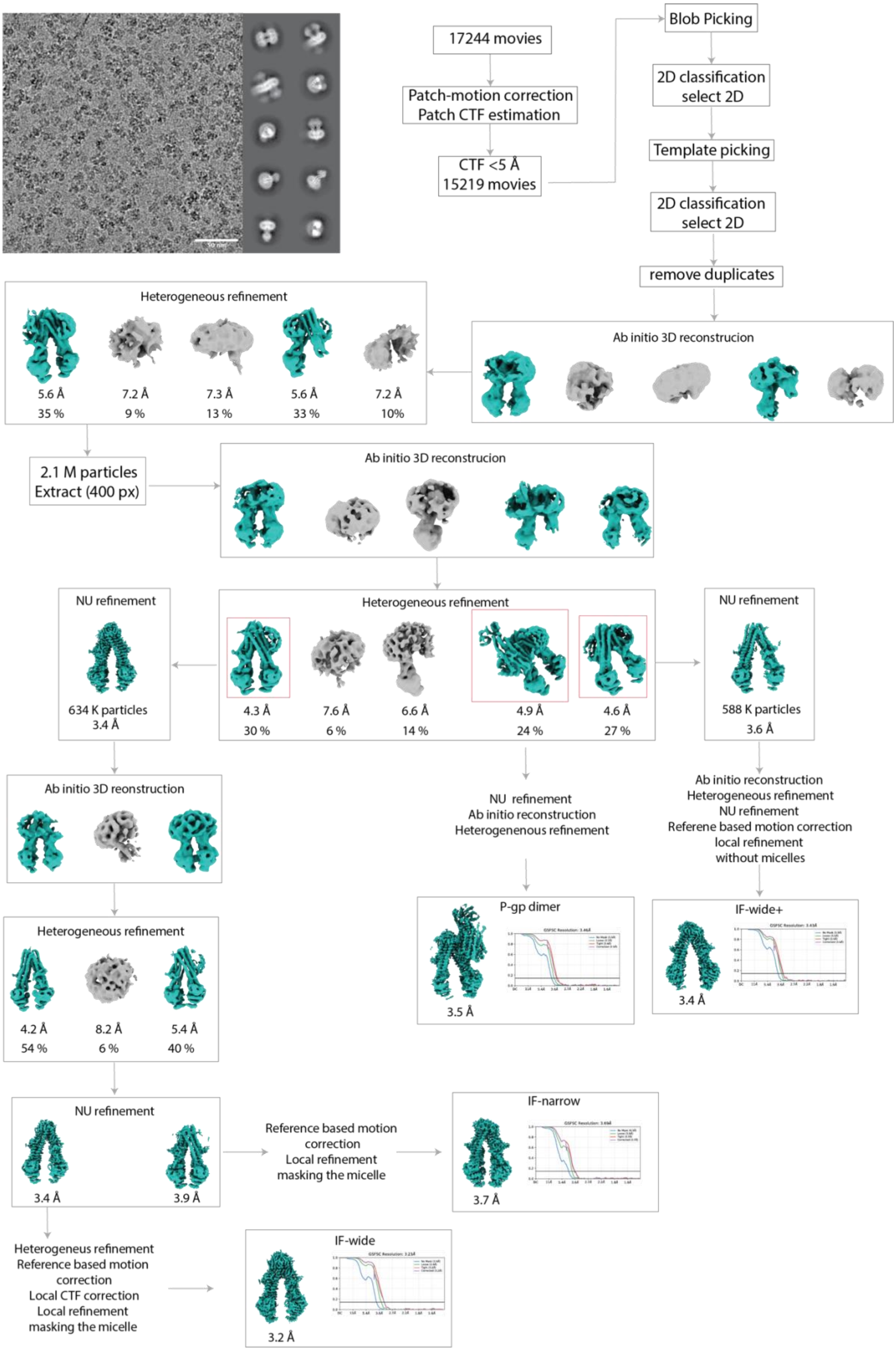

**Figure S8 Processing scheme of human P-gp in the apo state in detergent micelles.**

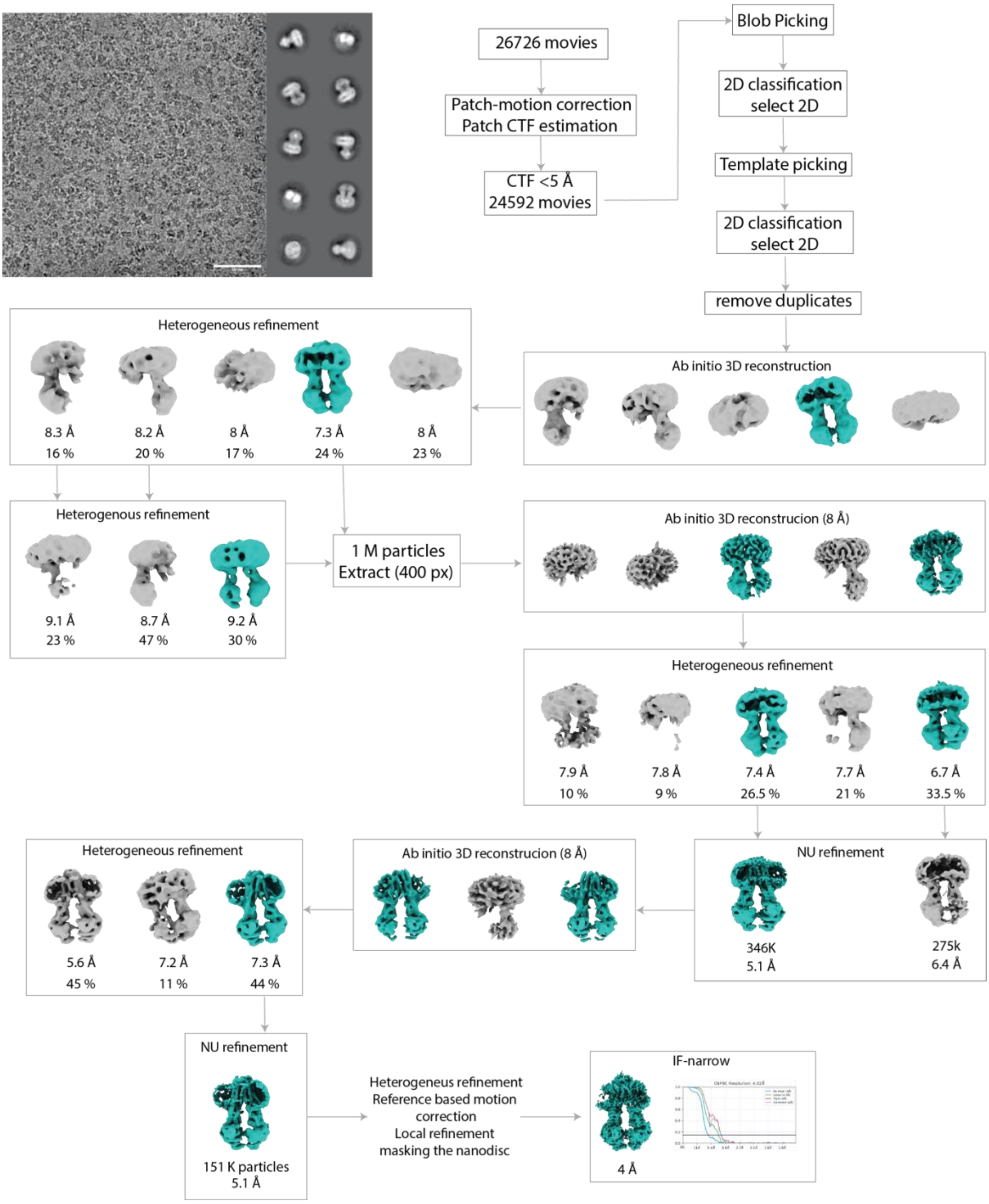

**Figure S9 Processing scheme of human P-gp in the verapamil-bound state in MSP2N2 nanodiscs.**

Human P-gp LMNG/CHS apo-wide+

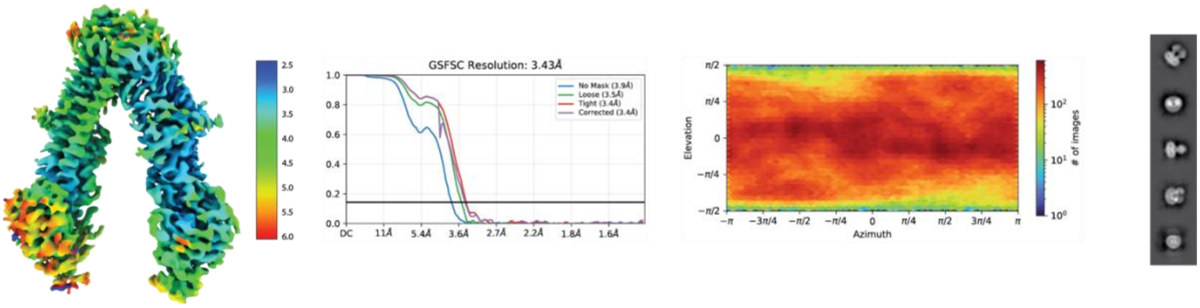

Human P-gp LMNG/CHS apo-wide

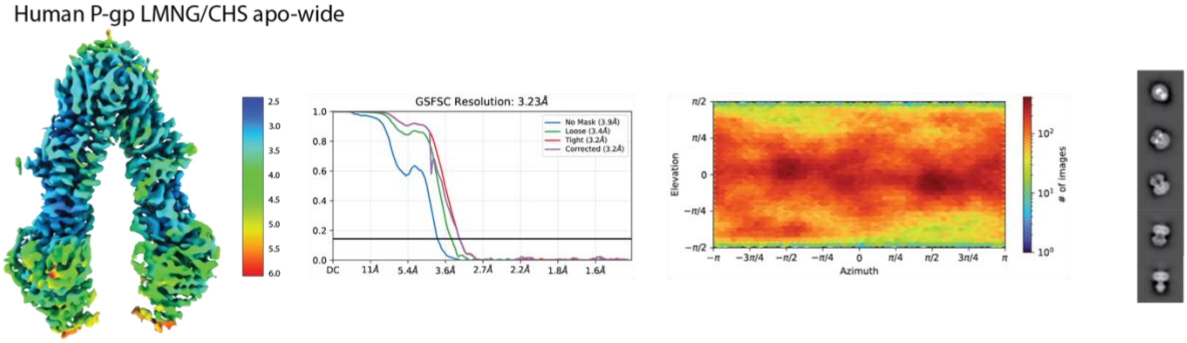

Human P-gp LMNG/CHS apo-narrow

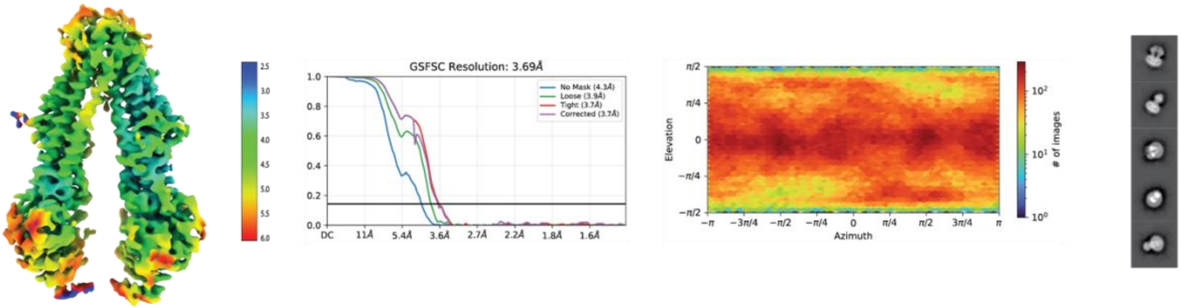

Human P-gp LMNG/CHS verapamil-wide

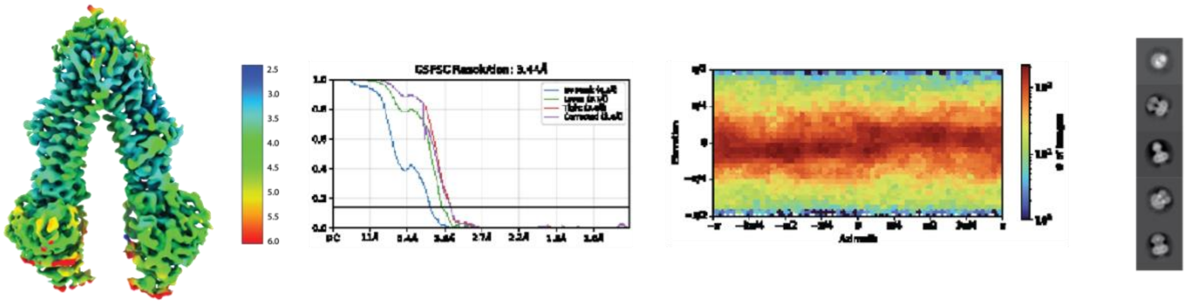

Human P-gp LMNG/CHS verapamil-narrow

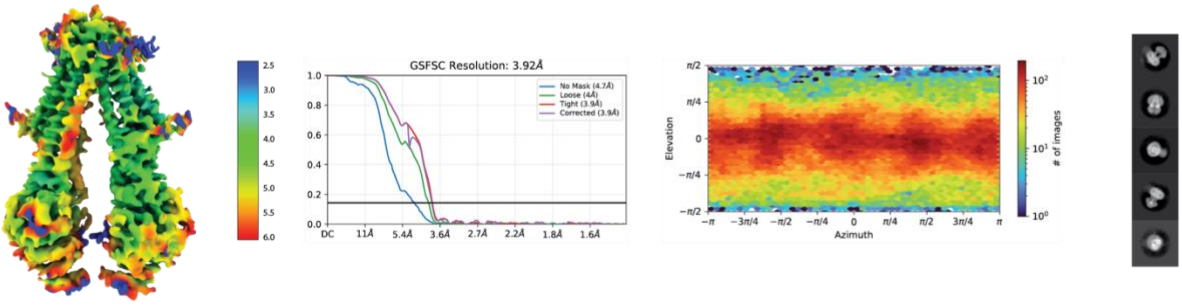

**Figure S10 Local resolution maps, FSCs, and orientation distribution plot for human P-gp in detergent micelles.**

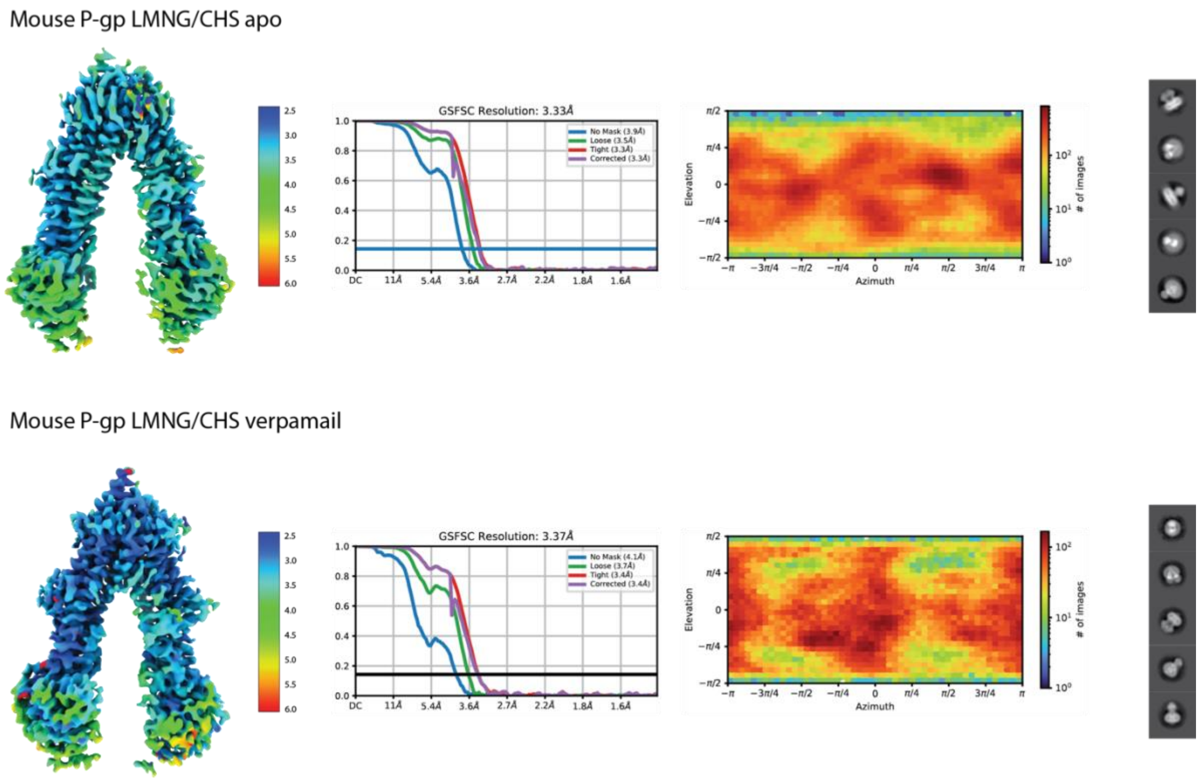

**Figure S11 Local resolution maps, FSCs, and orientation distribution plots for mouse P-gp in detergent micelles.**

Human P-gp MSP2N2 apo

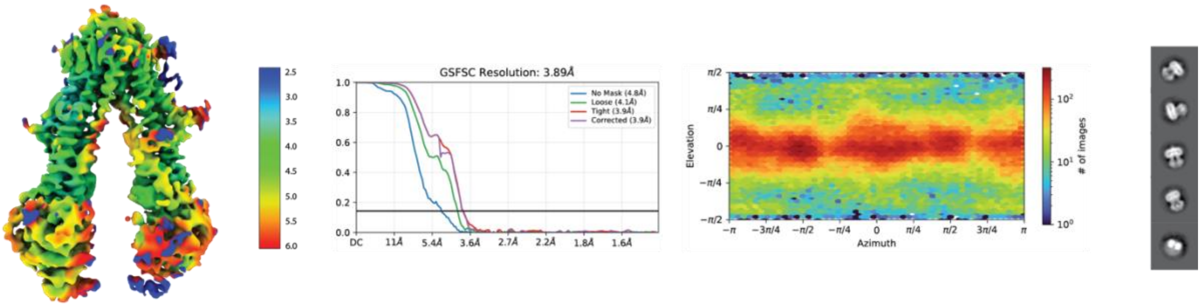

Human P-gp MSP2N2 verapamil

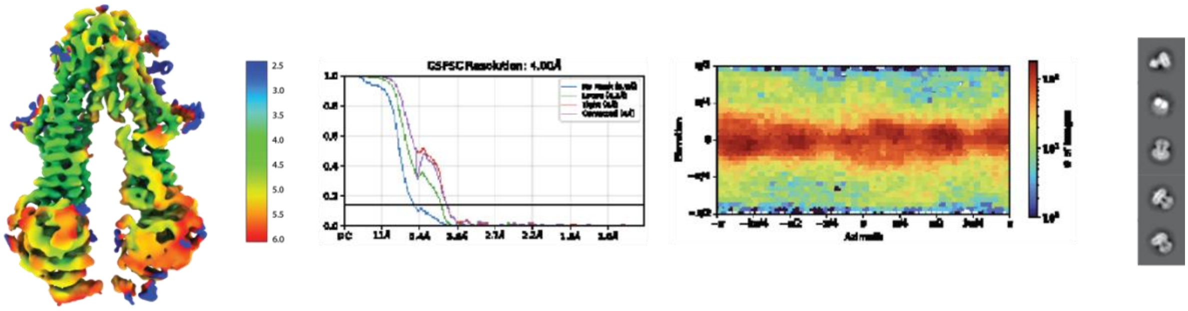

**Figure S12 Local resolution maps, FSCs, and orientation distribution plots for human P-gp in MSP2N2 nanodiscs.**

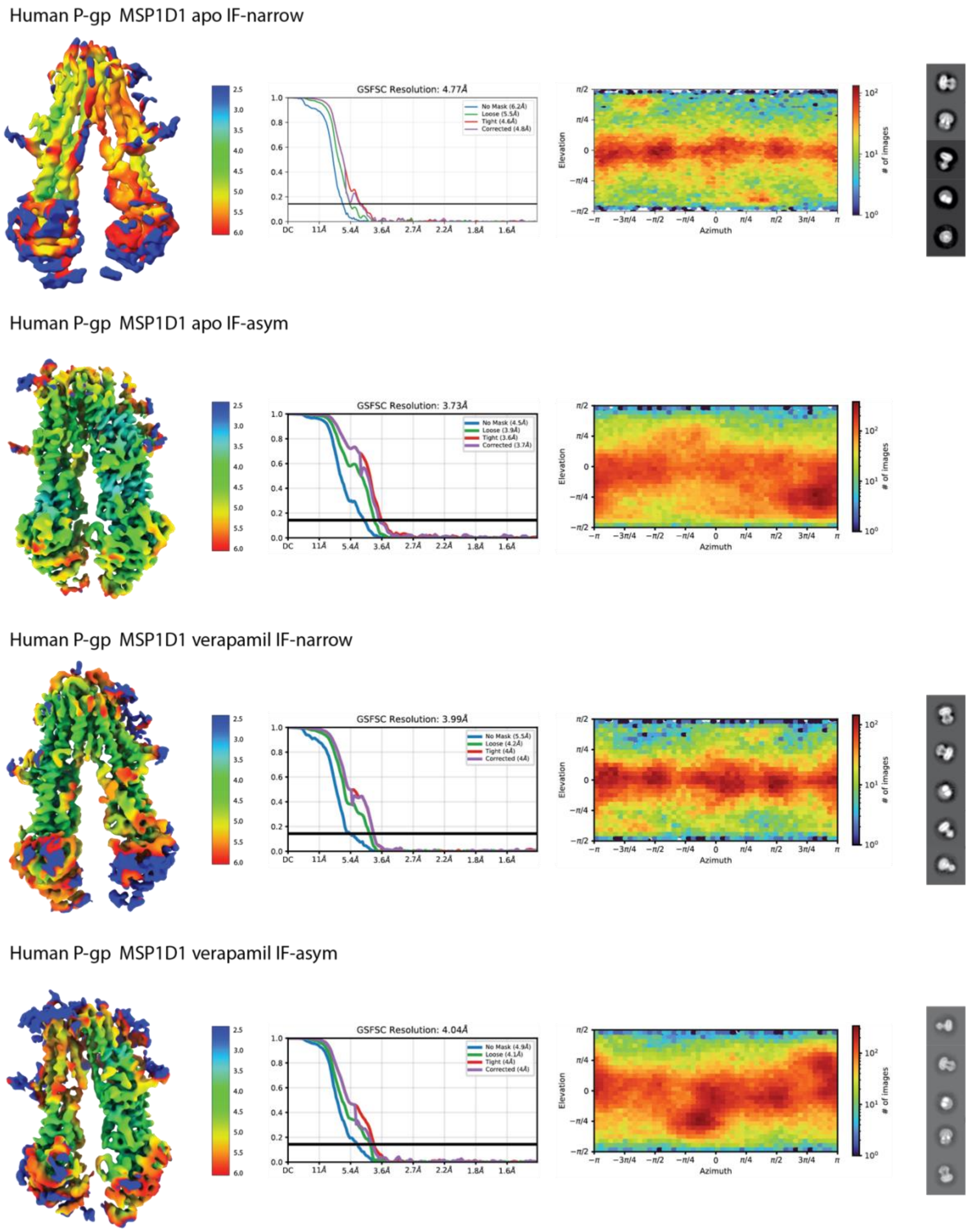

**Figure S13 Local resolution maps, FSCs, and orientation distribution plots for human P-gp in MSP1D1 nanodiscs.**

Table 1: **Cryo-EM data collection, refinement and validation statistics**

| | **hPgp-LMNGCHS-apo narrow** | **hPgp-LMNGCHS-apo wide** | **hPgp-LMNGCHS-apo wide +** | **hPgp-LMNGCHS-verapamil wide** | **hPgp-LMNGCHS-verapamil narrow** |
|---|---|---|---|---|---|
| **Data Collection** | | | | | |
| Accession number | EMD-56584 | EMD- 56585 | EMD- 56586 | EMD- 56587 | EMD- 56588 |
| Magnification | 165,000 | 165,000 | 165,000 | 165,000 | 165,000 |
| Voltage /kV | 200 | 200 | 200 | 200 | 200 |
| Dose / $e^{-}Å^{2}$ | 50 | 50 | 50 | 50 | 50 |
| Pixel size / Å | 0.68 | 0.68 | 0.68 | 0.68 | 0.68 |
| Defocus range / µm | -1.8 to -0.6 | -1.8 to -0.6 | -1.8 to -0.6 | -1.8 to -0.6 | -1.8 to -0.6 |
| Recorded movies | 17274 | 17274 | 17274 | 13810 | 13810 |
| Final particle images | 251585 | 322939 | 555974 | 156003 | 128845 |
| Microscope | Glacios | Glacios | Glacios | Glacios | Glacios |
| Camera | Falcon 4 | Falcon 4 | Falcon 4 | Falcon 4 | Falcon 4 |
| Energy Filter | Selectris | Selectris | Selectris | Selectris | Selectris |
| **Image Processing** | | | | | |
| Initial model | | | | | |
| Processing software | cryoSPARC (v.4) | cryoSPARC (v.4) | cryoSPARC (v.4) | cryoSPARC (v.4) | cryoSPARC (v.4) |
| Symmetry imposed | C1 | C1 | C1 | C1 | C1 |
| Resolution Å | 3.69 | 3.23 | 3.43 | 3.44 | 3.92 |
| Applied B-factor / Å2 | -149.3 | -118.2 | -117.4 | -98.3 | -118.1 |
| **Model Refinement** | | | | | |
| PDB accession | 28KU | 28KV | 28KX | 28KY | 28KZ |
| Validation | | | | | |
| FSCmap-to-model (0.143) / Å | | | | | |
| MolProbity score | 1.68 | 1.55 | 1.51 | 1.62 | 1.94 |
| Clash score | 3.74 | 3.65 | 2.47 | 3.88 | 7.26 |
| Composition | | | | | |
| Atoms | 9244 | 9667 | 9213 | 9188 | 9205 |
| Protein residues | 1184 | 1185 | 1185 | 1178 | 1180 |
| Ligands | - | Y01: 11 LMN: 1 | - | VER: 1 | VER: 1 |
| Bonds (R.M.S.D.) | | | | | |
| Length (Å) | 0.006 | 0.005 | 0.005 | 0.005 | 0.005 |
| Angles (°) | 0.987 | 0.895 | 0.800 | 0.803 | 0.923 |
| B-factors (min/max/mean) | | | | | |
| Protein residues | 38.34/149.76/89.17 | 15.74/162.36/76.03 | 18.00/194.80/92.39 | 34.67/182.25/96.50 | 70.32/233.19/144.64 |
| Ligand | - | 40.97/125.87/85.73 | - | 70.01/109.49/94.09 | 135.11/177.62/154.56 |
| Ramachandran plot (%) | | | | | |
| Favored | 91.17 | 94.15 | 93.21 | 93.09 | 90.36 |
| Allowed | 8.66 | 5.85 | 6.70 | 6.91 | 9.64 |
| Outliers | 0.17 | 0.00 | 0.08 | 0.00 | 0.00 |
| Rotamer outliers (%) | 0.51 | 0.61 | 1.12 | 0.21 | 0.51 |

| | hPgp-MSP1D1-apo-asym | hPgp-MSP1D1-apo-narrow | hPgp-MSP1D1-verapamil-narrow | hPgp-MSP1D1-verapamil-asym | hPgp-MSP2N2-apo-narrow |
|---|---|---|---|---|---|
| **Data Collection** | | | | | |
| Accession number | EMD- 56591 | EMD-56592 | EMD-56593 | EMD-56594 | EMD-56589 |
| Magnification | 165,000 | 165,000 | 165,000 | 165,000 | 165,000 |
| Voltage /kV | 200 | 200 | 200 | 200 | 200 |
| Dose / $e^{-}$Å$^{2}$ | 50 | 50 | 50 | 50 | 50 |
| Pixel size / Å | 0.68 | 0.68 | 0.68 | 0.68 | 0.68 |
| Defocus range / µm | -1.8 to -0.6 | -1.8 to -0.6 | -1.8 to -0.6 | -1.8 to -0.6 | -1.8 to -0.6 |
| Recorded movies | 18094 | 18094 | 28122 | 28122 | 41287 |
| Final particle images | 201439 | 59830 | 81048 | 200179 | 116314 |
| Microscope | Glacios | Glacios | Glacios | Glacios | Glacios |
| Camera | Falcon 4 | Falcon 4 | Falcon 4 | Falcon 4 | Falcon 4 |
| Energy Filter | Selectris | Selectris | Selectris | Selectris | Selectris |
| **Image Processing** | | | | | |
| Initial model | | | | | |
| Processing software | cryoSPARC (v.4) | cryoSPARC (v.4) | cryoSPARC (v.4) | cryoSPARC (v.4) | cryoSPARC (v.4) |
| Symmetry imposed | C1 | C1 | C1 | C1 | C1 |
| Resolution Å | 3.73 | 4.77 | 4.05 | 4.04 | 3.93 |
| Applied B-factor / Å2 | -133.9 | -261.8 | -103.5 | -137 | -134.7 |
| **Model Refinement** | | | | | |
| PDB accession | 28LC | 28LD | 28LE | XXXX | 28LA |
| Validation | | | | | |
| FSCmap-to-model (0.143) / Å | | X.X | | | |
| MolProbity score | 1.79 | 1.57 | 1.92 | | 2.01 |
| Clash score | 0.005 | 3.56 | 6.77 | | 8.43 |
| Composition | | | | | |
| Atoms | 8933 | 9159 | 9217 | | 9091 |
| Protein residues | 1149 | 1179 | 1180 | | 1168 |
| Ligands | - | - | VER: 1 | | |
| Bonds (R.M.S.D.) | | | | | |
| Length (Å) | 0.005 | 0.005 | 0.005 | | 0.006 |
| Angles (°) | 0.904 | 0.866 | 0.938 | | 1.035 |
| B-factors (min/max/mean) | | | | | |
| Protein residues | 66.33/160.08/111.25 | 30.00/326.82/229.52 | 118.82/273.67/184.32 | | 72.98/194.10/122.64 |
| Ligand | - | - | 158.24/158.24/158.24 | | |
| Ramachandran plot (%) | | | | | |
| Favored | 91.85 | 93.6 | 89.95 | | 89.85 |
| Allowed | 7.98 | 6.06 | 9.80 | | 9.55 |
| Outliers | 0.18 | 0.34 | 0.26 | | 0.60 |
| Rotamer outliers (%) | 0.95 | 0.10 | 0.51 | | 0.11 |

| | hPgp-MSP2N2-verapamil-narrow | mPgp-LMNGCHS-apo-wide + | mPgp-LMNGCHS-verapamil-wide |
|---|---|---|---|
| **Data Collection** | | | |
| Accession number | EMD- 56590 | EMD-56595 | EMD-56596 |
| Magnification | 165,000 | 165,000 | 165,000 |
| Voltage /kV | 200 | 200 | 200 |
| Dose / $e^{-}$Å$^{2}$ | 50 | 50 | 50 |
| Pixel size / Å | 0.68 | 0.68 | 0.68 |
| Defocus range / µm | -1.8 to -0.6 | -1.8 to -0.6 | -1.8 to -0.6 |
| Recorded movies | 26726 | 11039 | 12628 |
| Final particle images | 78890 | 450337 | 138286 |
| Microscope | Glacios | Glacios | Glacios |
| Camera | Falcon 4 | Falcon 4 | Falcon 4 |
| Energy Filter | Selectris | Selectris | Selectris |
| **Image Processing** | | | |
| Initial model | | | |
| Processing software | cryoSPARC (v.4) | cryoSPARC (v.4) | cryoSPARC (v.4) |
| Symmetry imposed | C1 | C1 | C1 |
| Resolution Å | 4.02 | 3.33 | 3.37 |
| Applied B-factor / Å2 | -136.6 | -131.4 | -99.4 |
| **Model Refinement** | | | |
| PDB accession | 28LB | 28LF | 28LG |
| Validation | | | |
| FSCmap-to-model (0.143) / Å | | X.X | |
| MolProbity score | 1.94 | 1.56 | 1.83 |
| Clash score | 7.57 | 2.87 | 9.25 |
| Composition | | | |
| Atoms | 9217 | 9386 | 9259 |
| Protein residues | 1180 | 1148 | 1131 |
| Ligands | VER: 1 | Y01: 14 | VER: 1, Y01: 9 LMN: 1 RVE: 1 3PE: 1 |
| Bonds (R.M.S.D.) | | | |
| Length (Å) | 0.006 | 0.005 | 0.005 |
| Angles (°) | 1.010 | 0.934 | 1.097 |
| B-factors (min/max/mean) | | | |
| Protein residues | 99.25/205.74/143.64 | 0.00/88.13.25.58 | 39.59/211.14/119.31 |
| Ligand | - | 8.08/42.48/19.62 | 59.17/155.02/109.02 |
| Ramachandran plot (%) | | | |
| Favored | 90.80 | 91.96 | 95.11 |
| Allowed | 9.20 | 7.95 | 4.89 |
| Outliers | 0.00 | 0.09 | 0.00 |
| Rotamer outliers (%) | 0.72 | 0.42 | 0.54 |
